# Native virion structure reveals stochastic genome assortment underlying dose-dependent Orsay virus infection

**DOI:** 10.64898/2026.07.28.741195

**Authors:** Soledad Stagnoli, Victoria G. Castiglioni, Lifei Fu, Ane Martinez-Castillo, Ander Vidaurrazaga, Susana Martín, Tammo Diercks, Santiago F. Elena, Nicola G.A. Abrescia

## Abstract

Orsay virus (OrV), infecting *Caenorhabditis elegans*, is a valuable model for studying host-virus interactions, yet its bipartite RNA genome packaging and assembly mechanisms remain unclear. To investigate these mechanisms, we combine infection assays, single-molecule imaging, and cryo-electron microscopy. We find that the two RNA segments accumulate asymmetrically, and that productive infection depends on coinfection by multiple particles. We therefore asked whether unequal segment availability generates heterogeneous virions. Cryo-EM analysis of OrV, including symmetric and asymmetric reconstructions to 2.5 Å resolution, unveils uniform internal density, excluding large populations of empty or semi-empty particles. We show the genome forms a semi-ordered network contacting the electropositive capsid interior through sequence-independent interactions with the N-terminus and penton region. We also resolve the capsid-8 linker, identifying the penton as a likely site coordinating capsid assembly. Together, we propose a model in which stochastic genome assortment emerges during assembly modulated by segment-specific replication dynamics.

## INTRODUCTION

Orsay virus (OrV) is the first natural viral pathogen identified in the nematode *Caenorhabditis elegans*. Its discovery opened a new avenue for studying host-virus interactions, as *C. elegans* serves as a model system for fundamental biological processes including development, aging, immunity, cellular dynamics, and gene regulation (*1*).

OrV is a non-enveloped, *T* = 3 icosahedral, positive-sense single-stranded RNA [(+)ssRNA] virus approximately 360 Å in diameter. Its bipartite genome is ∼6.0 kb long and consists of two RNA segments. RNA1 (∼3.4 kb) encodes the RNA-dependent RNA polymerase (RdRp), whereas RNA2 (∼2.6 kb) encodes the major capsid protein a, which contains a protruding domain (P) and a shell domain (S) (*2*). RNA2 also encodes 8, a protein required for viral egress (*3*). It has also been shown that a –1 ribosomal frameshift is required for translation of the extended a-8 fusion protein (∼84.5 kDa) and the 8 fiber protein is exposed on the virion surface (*4*). While not yet officially classified by the International Committee on Taxonomy of Viruses (ICTV; https://ictv.global/), OrV resembles members of the *Nodaviridae* family whose pathogenic members cause viral nervous necrosis in marine and freshwater fish, leading to significant aquaculture losses. Despite progress in understanding OrV infection and the tridimensional (3D) structure of virus-like particles (OrV-VLP) and their individual structural components, key determinants of genome encapsidation and virion assembly remain unknown (*2, 5, 6*).

For other RNA viruses, high-resolution structural studies have begun to decipher how RNA conformations and protein-RNA interactions orchestrate genome packaging. Studies of (+)ssRNA fiersvirus MS2 have demonstrated that its genome folds into defined tertiary structures that promote selective encapsidation through RNA packaging signals distributed along the genome (*7, 8*). Likewise, work on the cystovirus ϕ6 showed that its tri-segmented double-stranded RNA adopts a highly organized arrangement within the capsid, with coordinated RNA-protein interactions guiding sequential packaging and replication (*9*). More recently, analyses of human rhinovirus have revealed that its (+)ssRNA genome can adopt ordered organizations within the capsid, shaped by capsid-RNA contacts that stabilize the virion and regulate its uncoating (*10*). Together, these studies emphasize that decoding RNA organization inside native virions is essential for understanding the molecular mechanisms underlying genome packaging and virion assembly and disassembly.

Here, we investigated the segment packaging requirements for wild-type OrV infectivity and purified native virions from infected nematodes for high-resolution cryo-EM analysis. Single-molecule FISH (smFISH) experiments reveal asymmetric genome accumulation while dose-response assays indicate that productive replication often requires coinfection by multiple virions. Cryo-EM reconstructions of OrV at 2.5 Å resolution under icosahedral symmetry and ∼3.2 Å resolution under asymmetric or symmetry-relaxed conditions reveal interactions between the N-terminal domain of the capsid protein and the packaged RNA, forming a cage-like organization with annuli at the threefold and fivefold axes. Although the capsid could theoretically accommodate up to ∼3.6 RNA segments, geometric and electrostatic constraints likely limit this occupancy. Analysis of the internal average genome signal supports an average occupancy of ∼77% (*i.e*., ∼2.8 genome equivalents). Further, two-dimensional class averages of virions with masked capsids failed to reveal large virus populations with dramatic changes in genome content. We show that genome encapsidation is largely sequence-independent and primarily driven by the N-terminal region of the S domain which, in the presence of RNA, can assemble into nanoparticles resembling native virions in size, but lacking an ordered icosahedral lattice. Finally, we resolved the linker region of the a-8 fusion protein and its native fiber organization, establishing a structural basis for the role of –1 ribosomal frameshifting in the assembly of infectious particles.

Together, these findings indicate that OrV genome packaging is not strictly dependent on RNA segment identity, and we propose that replication dynamics, together with spatial confinement of RNA1 and RNA2, drive their co-assortment during assembly.

## RESULTS

### Coinfection by multiple virions is required for OrV productive replication

Previous studies have reported that *C. elegans* individuals may harbor OrV RNA1 even in the absence of RNA2 during infection (*11, 12*). To systematically investigate this phenomenon, we performed smFTSH on wild-type animals at early post-infection stages. At 8 hours post-infection (hpi), 82 ±2% of infected animals showed signals for both RNAs (Fig. 1a, b). However, 18 ±2% displayed signal only for RNA1 (paired-samples *t*-test on arcsine-square root transformed frequencies: *t*_2_ = 13.692, *P* = 0.005; effect size: *η*^2^ = 0.989), consistent with a temporal lag in RNA2 segment accumulation. This effect disappeared at 24 hpi when RNA2 was detectable in all infected individuals indicating that segment co-acquisition is ultimately achieved, but is not synchronized (Fig. 1a, b; two-samples *t*-test comparing RNA1-only frequencies at 8 and 24 hpi: *t*_4_ = 13.137, *P* < 0.001, *η*^2^ = 0.993).

**Figure 1.**
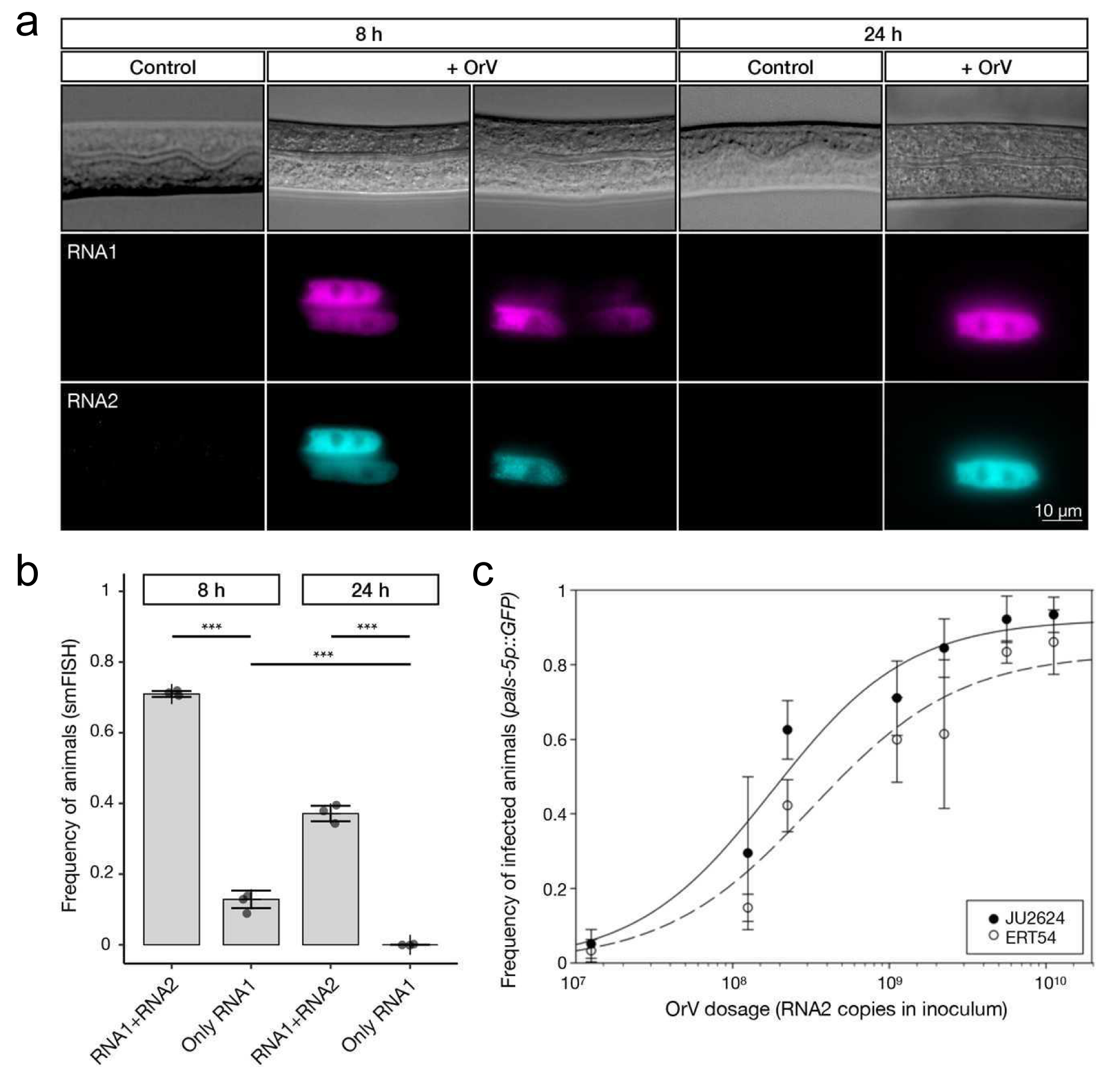
| Differential accumulation of OrV genomic segments and infection probability analysis. (**a**) Representative smFISH images of inoculated animals at 8 and 24 hpi. Top row: differential interference contrast (DIC); second row: RNA1 in magenta; third row: RNA2 in magenta. **(b)** Frequency of animals with RNA1 or RNA2 signals or only RNA1 signal determined by smFISH at 8 and 24 hpi; 20 to 24 animals per replicate, 3 biological replicates per time point. **(c)** Frequency of animals with activated GFP infection reporter at 24 hpi. 20 animals per replicate, 5 biological replicates per dilution. Bars indicate mean ± SD. Continuous lines represent the best fit to the dose-response model.

Since RNA1 encodes the RdRp, its earlier amplification could reflect delayed RNA2 synthesis or the existence of virions packaging RNA1 while lacking RNA2. To discriminate between these two scenarios, we tested the independent-action null hypothesis (IAH) (*13, 14*). IAH is the simplest model of virus infection kinetics and assumes that each virion has a non-zero probability to infect a cell (*ρ*) independent of the inoculation dose (*D*), and that virions act independently of one another (*i.e*., in a Poisson process) throughout the infection process. IAH can be tested by data fitting to the expression I(D) = 1 – exp(–pD^K^), where *I*(*D*) is the number of infected individuals for a given dose *D* of virions. If the IAH model holds, power *K* = 1 and infection is a purely uncorrelated Poisson process with success rate *ρ*. If *K* ≠ 1, infection becomes dose-dependent. If not all virions are infectious, *K* < 1 and *D* reduces to a so-called effective dose; *i.e.*, successful infections require a disproportionally large amount of inoculum. For example, for bipartite plant viruses it has been shown that *K* = 0.5 and the effective dose scales with the square root of the total dose (*15*).

We performed OrV dose-response experiments in wild-type ERT54 and hypersusceptible JU2624 strains of *C.elegans* using transcriptional GFP reporters of infection (*pals-5p::GFP* or *lys-3p::GFP*, respectively). Fig 1c shows the results of the experiments, revealing that the data can be well fitted to the cited *I*(*D*) equation (*R*^2^ = 0.908, *F*_4,76_ = 591,091, *P* < 0.001). As expected, the hypersusceptible JU2624 strain showed ∼2-fold higher infection success rates (*ρ* = 4.102 ±0.708) than ERT54 (*ρ =* 2.304 ±0.759). However, both strains yielded the same estimated *K* = 0.478 ±0.051. As with bipartite plant viruses, this reduced power *K* ≈ 0.5 indicates that the effective infectious dose is lower than expected, and that either two particles are required for a productive infection or that only some 50% of all particles are infectious.

### Overall cryo-EM structure of OrV

To assess whether differences in genome segment packaging could explain the two-fold higher dose required to achieve productive infection, we purified virions and determined the icosahedral structure of OrV by single-particle cryo-EM at 2.5 Å resolution using RELION-v4 (*16*) (Fig. 2a, left; Supplementary Figs. 1 and 2a). In addition, we performed 3D refinements using an expanded particle dataset with and without imposed icosahedral symmetry (Supplementary Table 1 and Supplementary Fig. 2b). For the latter, refinements included focused classification and symmetry-relaxation procedures implemented in cryoSPARC v4.7 (*17*). The capsid measures approximately 360 Å from vertex to vertex, consistent with the previously reported OrV-VLP model (*2*). Unlike the VLP structure, however, the native OrV reconstruction displays additional density beneath the capsid shell corresponding to the packaged RNA genome, as well as fiber-like spikes located at the fivefold symmetry axes (Fig. 2a right, and Fig. 2b). Docking of the previously reported 3.25 Å X-ray structure (PDB code 4NWV, https://doi.org/10.2210/pdb4NWV/pdb) of the three quasi-equivalent capsid proteins that form the icosahedral asymmetric unit (IAU), followed by real-space refinement, resulted in minor backbone rearrangements (root-mean-square deviation ≤ 0.54 Å). This procedure, however, yielded an atomic model of the IAU with substantially improved stereochemical quality (Fig. 2c and Supplementary Table 1). All 180 capsid subunits adopt a canonical jelly-roll B-barrel fold in the shell (S) domain and contain the insertion forming the B-rich protruding (P) domain, although the corresponding densities display variable levels of order as revealed by local resolution estimation (Fig. 2c and Supplementary Fig. 3a). The three quasi-equivalent subunits (A, B, and C; right middle inset in Fig. 2c) display differences in their terminal regions that reflect distinct local environments at the two-, three-, and five-fold symmetry axes (Supplementary Fig. 3b). The density corresponding to subunit A extends further toward the N-terminus, with Ser30 as the first conservatively modelled residue, compared with Asn46 and Ser44 in subunits B and C, respectively. Low-pass filtering reveals additional weaker density suggestive of a short helix that merges into a bulkier density located beneath the capsid at the threefold symmetry axis (Supplementary Fig. 3c). AlphaFold3 (*18*) predicts that the remaining twenty-nine N-terminal residues form a slightly bent a-helix (Fig. 2c, bottom inset). This N-terminal domain is highly positively charged and at the icosahedral threefold axes converges to form a positively charged cluster, together with the overall electropositive character of the internal capsid surface, inclusive of the fivefold vertex (Fig. 2d, e and Supplementary Fig. 4). The unresolved N-terminal ∼44 residues in the B and C subunits is consistent with structural flexibility. In contrast, the C-terminal extension of subunit A projects toward the fivefold vertices, forming a platform from which the 8 fibers emerge (Fig. 2b). Tn subunits B and C, the final modelled C-terminal residues are Pro388 and Ser389, respectively, and are located near the twofold icosahedral axis.

**Figure 2.**
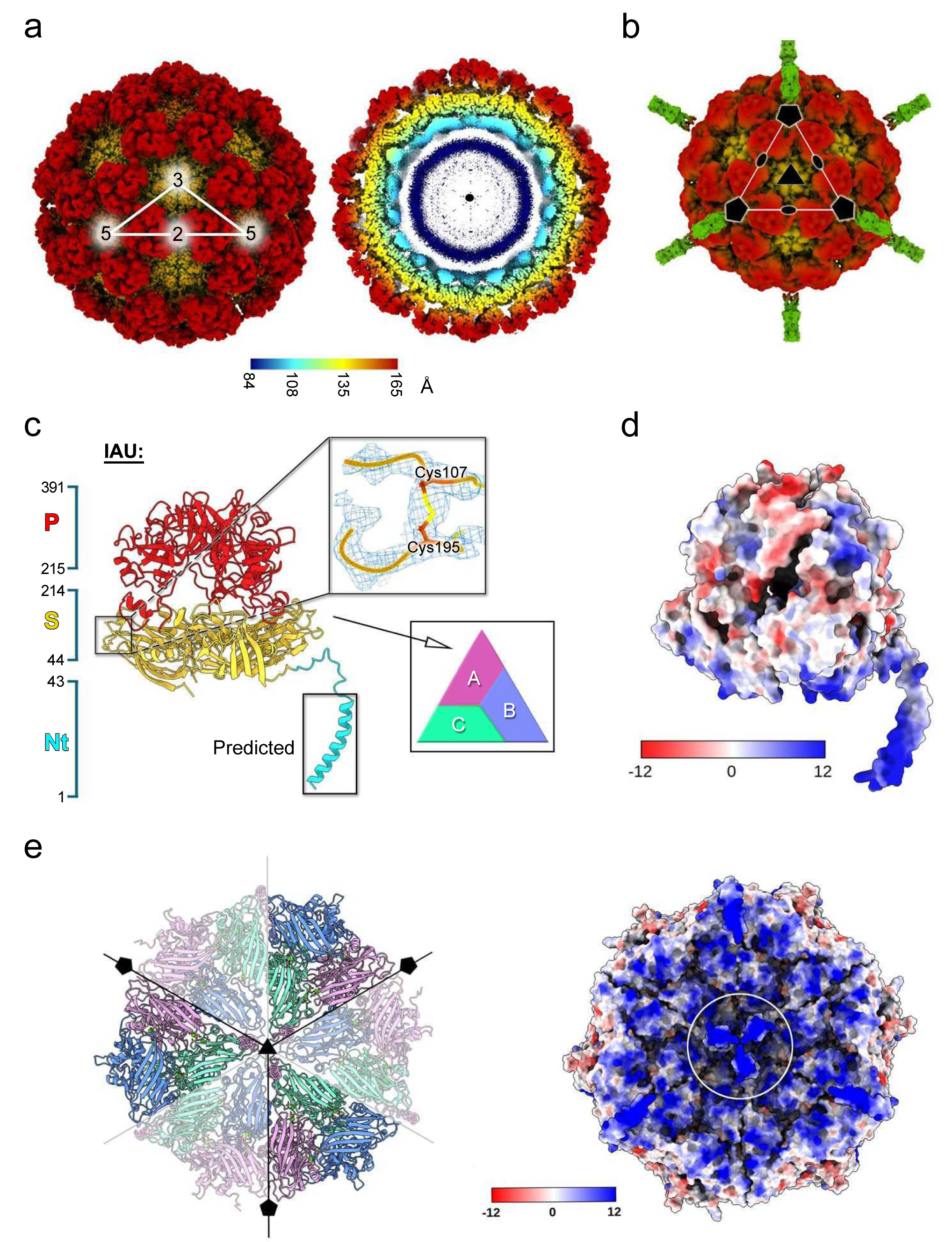
| Overall cryo-EM structure of the wt-OrV capsid. (**a**) Left, surface rendering of the cryo-EM icosahedral density map of the OrV virion at 2.5 Å resolution, showing the ∼360 Å diameter capsid. The white triangle indicates the icosahedral asymmetric unit (IAU), and the numbers 2, 3, and 5 denote the icosahedral two-, three-, and five-fold symmetry axes. Right, cutaway view showing internal density corresponding to the packaged bipartite RNA genome, color-coded by radius according to the key. **(b)** Surface rendering displayed at lower threshold showing protruding fibers at the fivefold vertices. A virus facet is outlined in white, and pentagon, oval, and triangle symbols indicate the icosahedral symmetry axes. **(c)** The IAU contains three quasi-equivalent a capsid proteins (A, B, and C; bottom inset), each composed of a shell (S, yellow) and protruding (P, red) domain adopting a jelly-roll B-barrel fold; domain boundaries are indicated on the left. Subunit A contains additional resolved N-terminal residues up to Ser30, whereas the preceding residues are predicted to form an a-helix extending beneath the threefold axis. Top inset, detailed density of subunit B (or C) showing the Cys107-Cys195 intradomain disulfide bond. (**d**) Electrostatic surface representation of the IAU viewed as in (c). (**e**) Left, bottom view of IAUs converging at the threefold axis (black triangle), with adjacent IAUs and N-terminal helices visible at the center (light magenta); black pentagon symbols indicate the fivefold axes. Right, corresponding electrostatic surface representation. The white circle highlights the highly electropositive N-terminal helices converging beneath the threefold axis.

Intermolecular interactions within the IAU vary depending on subunit position. Salt bridges and hydrogen bonds are observed across all interfaces, where hydrogen bonding is more extensive at the A-C interface, consistent with a higher propensity to form A-C *vs* A-B and B-C dimers (interface areas: A-C = 1772.8 Å^2^; B-C = 1471.6 Å^2^; A-B = 1449.4 Å^2^) (Supplementary Fig. 3b). Three Ca2+ ions bridge each interface between adjacent S domains and are coordinated by residues Glu166 and Gln92 from chain A, and Asp122 and Asp125 from chain C, as previously observed in the crystallographic virus-like particle structure (*2*). In addition, two further water molecules were identified coordinating the Ca2+ ions (Supplementary Fig. 3d). A total of 210 water molecules were modelled per IAU, where approximately 85% are located in proximity to the S domain. We attribute this asymmetric water distribution to the increased flexibility of the P domain (Supplementary Fig. 3a). Consistently, weaker density is observed for the exposed loops encompassing its residues Val305-Gly310 in all three subunits, indicative of enhanced local conformational mobility.

Unlike in the VLP crystal structure, disulfide bonds between Cys107 and Cys195 are observed in the S domain of subunits B and C, but are absent in subunit A (Fig. 2c, inset; Supplementary Fig. 3e). Cys107 directly precedes the Thr109-Thr111 loop, which, in the case of subunit A, converges at the fivefold axis to form a plateau-like structure that can accommodate the 8 fiber protein (see below) (Supplementary Fig. 3f).

### Genome organization and capsid interactions

Cryo-EM reconstructions of OrV, calculated from either a restricted particle set refined with icosahedral symmetry or an expanded particle set refined with relaxed or no imposed symmetry, revealed similar internal densities corresponding to the encapsidated RNA genome (Fig. 3a, b and Supplementary Fig. 5a). This similarity may reflect either the absence of a consistent genome organization across particles or the dominance of the highly symmetric capsid signal, which may obscure density arising from a potentially ordered RNA genome. Because ssRNA1 and ssRNA2 are inherently asymmetric molecules that are predicted by RNAfold (*19*) to adopt complex secondary structural motifs, including stem-loops and bulges (Supplementary Fig. 5b), the resulting symmetry-averaged genome density cannot be precisely interpreted. Nevertheless, several structural features are consistently observed in both unsharpened symmetrical (restricted and expanded dataset) and non-symmetrical reconstructions. The genome-associated density is organized into two distinct regions. The first comprises a relatively strong, semi-ordered cage-like lattice immediately beneath the inner capsid surface, with prominent densities located beneath the icosahedral threefold and fivefold axes (Figs. 2a right and 3a). Density beneath the threefold axes is interconnected by branching densities related by twofold symmetry, which extend toward and merge with a bell-shaped density beneath the fivefold axes (Fig. 3b and Supplementary Fig. 5a). A second, weaker genome-associated density forms a diffuse ∼30 Å-thick spherical shell centered ∼90 Å from the particle center, with density progressively fading toward the core. This organization is indeed captured by the radial density profile of the icosahedral OrV reconstruction, which we compared with profiles derived from OrV-VLPs and from a simulated density map calculated solely from the capsid atomic model, serving as an empty-particle control (Fig. 3c). After normalization to the intensity of the empty capsid shell, the OrV profile showed two distinct shoulders beneath the inner capsid surface, with relative intensities of ∼60% and ∼23%, respectively. The higher-intensity shoulder, located between radii of 90 and 110 Å, corresponds to the semi-ordered genome layer immediately beneath the capsid and merges with the inner boundary of the capsid density. The lower-intensity shoulder, spanning radii of 60-90 Å, corresponds to the deeper, more diffuse genome-associated density. Below a radius of 60 Å, the profile becomes largely featureless, indicating that the genome adopts a progressively less ordered organization toward the particle center. In contrast, the empty-particle control displays the expected flat profile beneath the capsid shell, whereas the OrV-VLP reconstruction exhibits only a single weak shoulder (∼20% normalized intensity), consistent with previous reports describing incorporation of short, nonspecific RNAs (<200 nucleotides) during recombinant capsid protein expression in *Escherichia coli* cells (*2*) (Fig. 3c).

**Figure 3.**
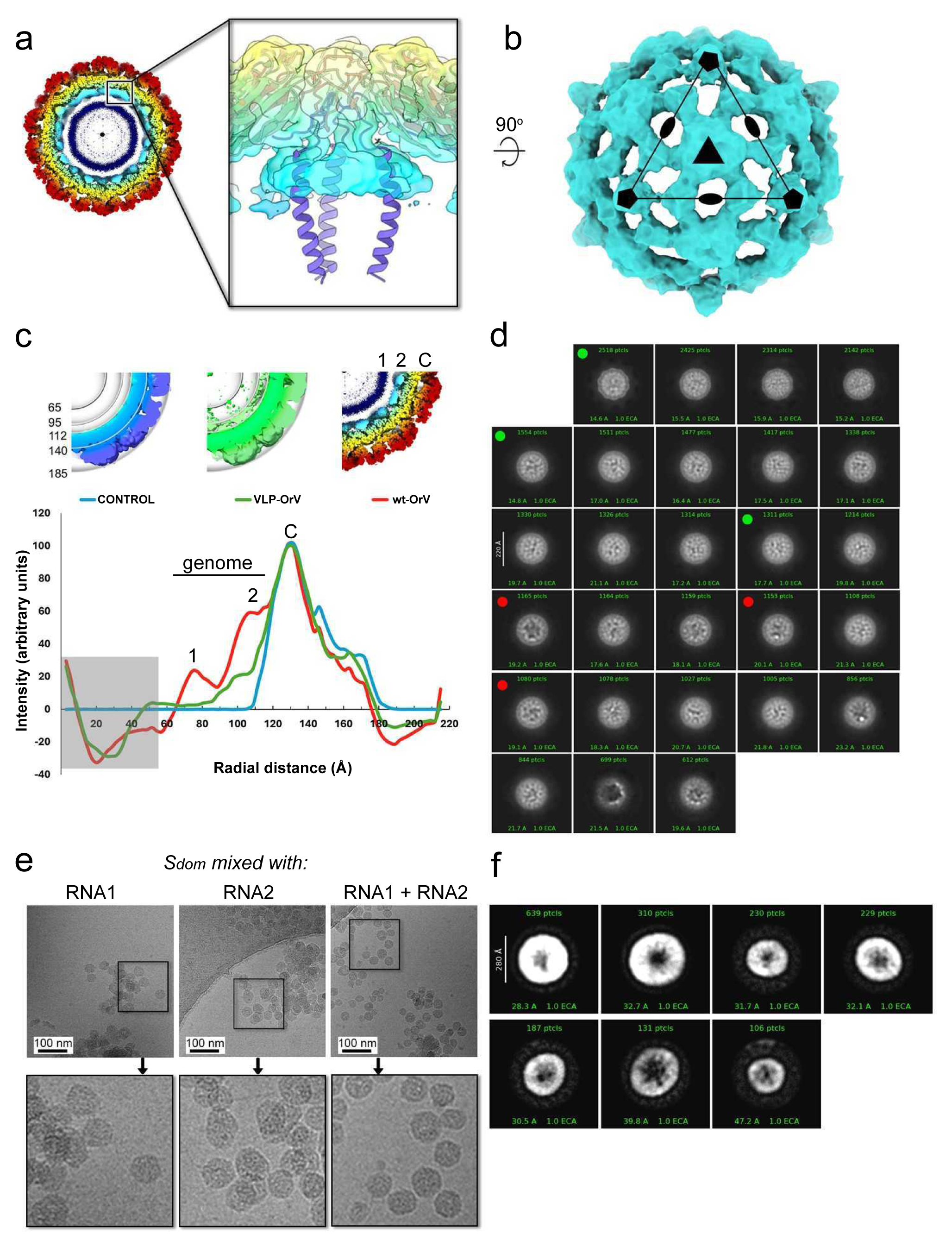
| RNA-protein anchoring network and genome content variability in OrV particles. (**a**) Cutaway view of the native OrV icosahedral reconstruction, with an enlarged inset showing the semi-transparent virion density. Fitted N-terminal predicted a-helices of subunit A (light purple) form protein-RNA clamps at the threefold axes, tethering the genome to the capsid shell. (**b**) Icosahedrally averaged genome density (cyan; low-pass filtered to 10 Å), revealing a cage-like network of density beneath the symmetry axes. (**c**) Radial density profiles of the control particle, OrV-VLP, and native OrV, showing the distribution of internal genome density in OrV (regions 1 and 2) and its close association with the inner surface of the capsid shell (C). The grey shaded region below ∼55 Å denotes the excluded central volume, which exhibits only diffuse density. (**d**) 2D classification of OrV particles following masking of the capsid shell and classification using only the internal density corresponding to the genome. Green circles (top left of class averages) highlight representative classes of genome-filled particles, whereas red circles indicate classes displaying reduced internal density, suggesting lower genome content relative to filled particles. Most classes displayed comparable internal density corresponding to genome-filled particles. (**e**) 2D cryo-EM images of nanoparticles assembled by incubating the S domain (14.8 µM) with the corresponding viral genome segments (RNA1, RNA2, or RNA1 + RNA2; 0.2 µM total RNA). Representative micrographs are shown at the top, with boxed regions enlarged threefold below to highlight particle morphology. Scale bars, 100 nm. (**f**) Representative 2D class averages of the assembled nanoparticles, illustrating structural heterogeneity in particle size while retaining common architectural features, including a high-contrast peripheral density and a central cavity visible as a darker region. The number of particles contributing to each class and the corresponding estimated resolution are indicated.

Electrostatic surface analysis highlights a marked contrast between the capsid exterior and interior (Fig. 2d, e, Supplementary Figs 4). Whereas the external capsid surface is largely neutral to negatively charged, the interior surface is strongly and nearly continuously electropositive. At each threefold axis, three putative N-terminal helices from the A capsid subunits converge to form positively charged protrusions enriched in Lys and Arg residues (positions 2, 7, 10, 11, 13, 16, 22, 23, 29, 36, and 41), that likely serve as anchors for the negatively charged phosphate backbone of the RNA genome or hydrogen bonds with bases (Fig. 3a, and Supplementary Figs. 3c and 4a). Although the electropositive-rich N-terminal regions of the B and C subunits are not resolved, they may similarly contribute to asymmetric genome organization within the capsid interior. At the fivefold axis, on the interior surface of the peripentonal capsid proteins, a ring of five Lys150 residues defines a ∼12 Å diameter crown that generates a localized region of positive electrostatic potential. This region extends into a broader positively charged surface patch comprising residues starting at Lys100 and Arg102, which coincides spatially with the larger section of the bell-shaped genome density observed beneath the fivefold axes (Supplementary Fig. 4b). To explore the nature of interactions between N-terminal domain of the capsid protein and vicinal genome at the threefold axis, we employed AlphaFold3 to generate a protein-RNA interface model (*18*). While inherently speculative, the model positions the N-terminal arm (blue helix; Fig. 2d) such that basic side chains (Lys, Arg and His) are oriented towards the RNA phosphate backbone to form an extended network of putative intermolecular hydrogen bonds and salt bridges. Additional polar interactions involving Asn and Ser residues may further stabilize the local geometry (Supplementary Fig. 6). To estimate genome occupancy, we then compared the total volume enclosed by the capsid (∼6.0×10^6^ Å^3^) with the icosahedrally averaged genome-associated density contoured at 0.0025a in ChimeraX (*20*). At this threshold, the RNA occupies ∼4.6×10^6^ Å^3^, indicating that at least 75% of the internal capsid volume is filled. Given that the theoretical volume of a single copy of RNA1 and RNA2 is about 1.65×10^6^ Å^3^, this suggests that the capsid can, in principle, accommodate up to ∼3.6 genome segments. To assess variation in genome content among virions, we performed focused 2D classification of internal densities from the large dataset of particles contributing to the OrV 3D reconstruction, as conventional 2D classification of extracted virions did not reveal any classes consistent with empty or partially filled particles (Supplementary Fig. 2b, top right). After masking the capsid shell, we identified a subset of particles displaying distinct contrast within the genome region, corresponding to an estimated ∼8% of the (71,626) classified particles (Fig. 3d, red dots). Further 3D classification of this subset did not uncover reproducible structural differences.

**Figure 4.**
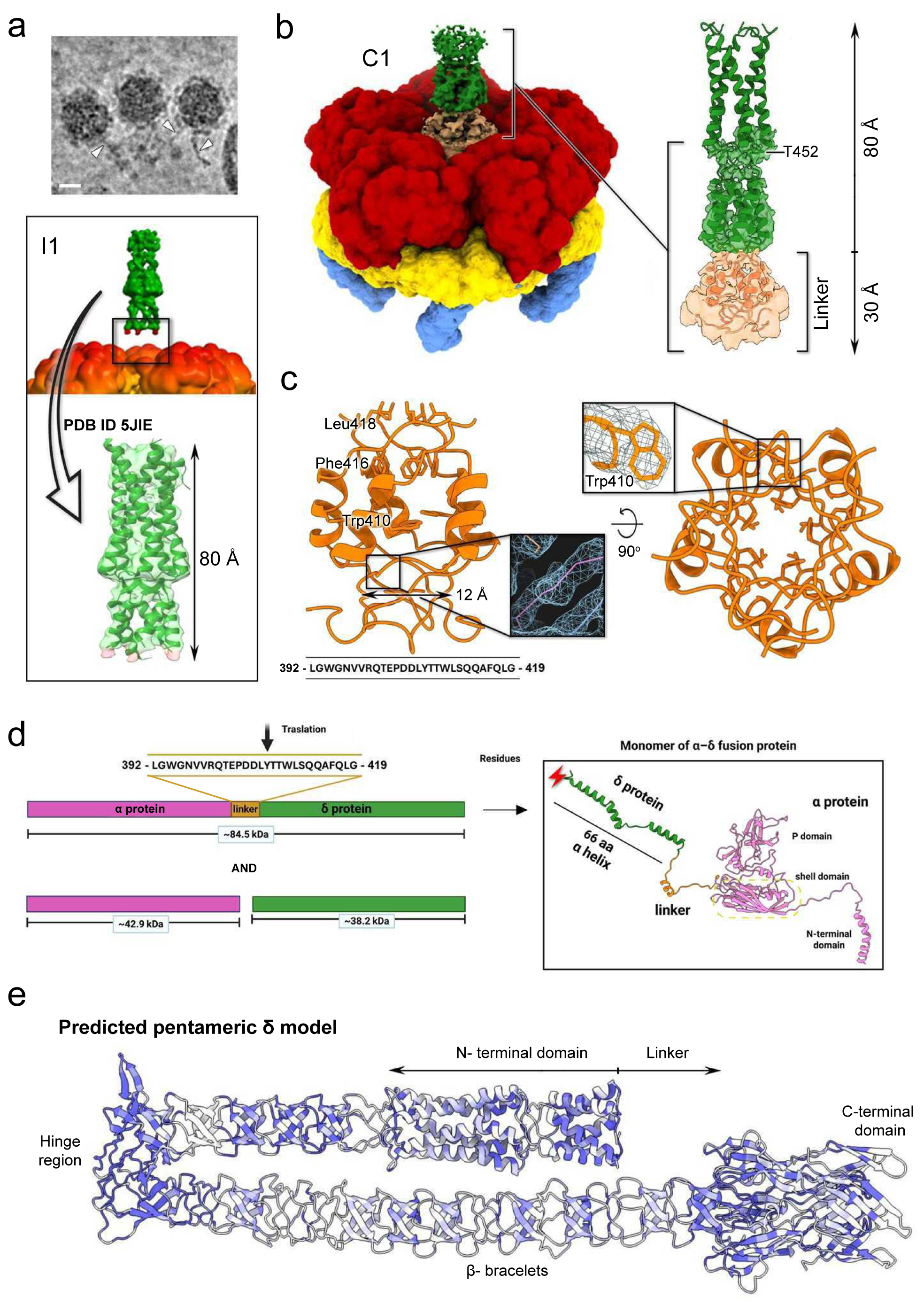
I Structural organization and capsid anchoring of the 6 fiber. (**a**) Top, cryo-EM micrograph showing 8 fibers (white arrowheads) projecting from fivefold vertices; scale bar, 250 Å. Bottom, icosahedrally averaged capsid reconstruction (red/yellow) showing additional density (green) corresponding to a fragment of the 8 fiber extending from the vertices, but without continuous density attached to the capsid. The green density is fitted with the available crystal structure (PDB ID 5JIE; black-outlined arrow). (**b**) Localized asymmetric (C1) 3D refinement (color-coded as in Fig. 2c) resolves the N-terminal stalk of the linker connecting the capsid and 8 fiber (-30 A linker height), extending from the C terminus of the a capsid subunit A (right inset). (**c**) Left, side view of the resolved linker region, showing a coiled-coil with a hydrophobic core connecting the a capsid to the 8 fiber docking interface (inset shows corresponding density in light blue visualized in Coot, with backbone shown in stick representation). Rings of Phe416 and Leu418 orient toward the fiber interior. Right, bottom view of the linker, with an inset highlighting density surrounding residue Trp410. Insets are not shown to scale. (**d**) Left, schematic representation of the concatenated structural proteins, with the linker positioned between domains. Ribosomal frameshifting generates a full-length a-8 fusion protein (top), whereas in its absence a and 8 are produced as separate proteins (bottom). Right, schematic of the full-length a-8 protein shown as a cartoon, color-coded as in the left panel; red lightning symbol indicates that this is not the full-length 8 protein. (**e**) AlphaFold3-predicted pentameric 8 fiber, colored by sequence conservation across Le Blanc and Santeuil viruses. Color scale: dark blue, strictly conserved residues; light blue, moderately conserved; white, low identity or gaps (variable regions).

### N-terminal arm– and RNA-dependent self-assembly of S-domain nanoparticles with sequence-independent encapsidation

We next investigated whether the S domain alone (S_dom_) of the a protein was sufficient to promote genome encapsidation and particle formation, given that the full-length capsid protein has previously been shown to self-assemble into practically empty OrV-VLP (*2*). Purified S_dom_ was predominantly monomeric in solution, although it showed a propensity to form oligomers (Supplementary Fig. 7a-b). This was further supported by nuclear magnetic resonance (NMR) spectroscopy, where increasing S_dom_ concentrations, led to progressively signal broadening and intensity loss (up to ∼20% reduction at 130 µM *vs* 10 µM protein concentration), together with a decrease in the apparent diffusion coefficient (Supplementary Fig. 7c-d). This is in line with some concentration dependent self-association of the S_dom_, although larger aggregates (> 100 kDa) are not resolvable by direct NMR observation. Upon addition of RNA1 and RNA2 (0.2 µM total) in a 130 µM S_dom_ solution, the 1D ^1^H NMR spectrum exhibited an additional ∼20% reduction in signal intensity. This decrease is consistent with partial RNA-induced formation of higher-order supramolecular assemblies that are not detectable by NMR due to their large molecular size (Supplementary Fig. 7d). Consistent with these findings, incubation of 14.8 µM S_dom_ with 0.2 µM viral RNA1, RNA2, or a mixture of both resulted in the formation of nanoparticles as visualized by 2D cryo-EM (Fig. 3e). Prior to this, to assess whether the two RNA segments interact with each other, we performed an electrophoretic mobility shift assay. Whether mixed immediately before loading or pre-annealed prior to electrophoresis, the two RNA segments migrated as distinct bands, providing no evidence of stable intermolecular interactions under the conditions tested (Supplementary Fig. 8). Nanoparticle assembly was independent of the RNA segment used; however, the formation of spherical particles appeared to require a minimum RNA concentration of 0.2 µM in the presence of 14.8 µM S_dom_. Two-dimensional classification of 1,832 particles assembled in the presence of both RNA1 and RNA2 revealed smooth, spherical structures with diameters ranging from ∼190 to ∼280 Å. These particles lacked apparent icosahedral symmetry, with the most populated classes exhibiting diameters of ∼280 Å (Fig. 3F), comparable to the diameter of native OrV excluding the protruding P domain. Notably, the nanoparticles displayed a variable radial electron density distribution, with a low-density core and increased density toward the inner surface of the shell, reminiscent of the genome-containing region observed in a subset of OrV 2D class averages and consistent with RNA localization along the interior of the S_dom_ shell (Fig. 3f). This distribution mirrors the electrostatically mediated genome organization observed in the OrV 3D density maps (Fig. 3c, d).

To examine RNA sequence dependence, we next tested nanoparticle assembly in the presence of heterologous nucleic acids. Unrelated viral ssRNA (6 kb) from turnip mosaic virus (TuMV) promoted S_dom_ assembly under equivalent RNA1/2:S_dom_ ratios, but yielded larger and more heterogeneous nanoparticles that formed even at lower protein concentrations (≥ 0.1 µM) (Supplementary Fig. 9a). In contrast, short ssDNA fragments (39, 50 and 100 nucleotides) predicted to form bubbles and hairpins (*19*), failed to produce co-assembled structures as by cryo-EM, despite binding to the S_dom_ as assessed by electrophoretic mobility shift assay (EMSA) analysis (Supplementary Fig. 9b). By comparison, longer linear DNA (∼2 kb), also predicted to form secondary structures, restored particle formation and generated a heterogeneous population of spherical and tubular particles (Supplementary Fig. 9c). Thus, nucleotide length further influences assembly outcomes, and relatively long ssDNA substrates can lead to mixed morphologies.

Finally, to assess the role of the first forty N-terminal in genome encapsidation and nanoparticle assembly, we generated a deletion mutant lacking the N-terminal arm (ilN-S_dom_, residues 42-211). In contrast to S_dom_, ilN-S_dom_ failed to assemble into detectable nanoparticles under identical conditions with RNA1, RNA2, or their combination (Supplementary Fig. 10).

### Incorporation of the 6 fiber surface protein into the virion

We observed protruding fibers in a subset of OrV particles in raw cryo-EM images (Fig. 4a, top). Consistently, the icosahedrally averaged 3D reconstruction showed additional density at all fivefold vertices, although at lower threshold than the capsid. However, we did not detect continuous density connecting the fibers to the capsid shell, nor density corresponding to the full-length 8 protein (residues Met420-Asp765 in a-8 fusion protein) (Figs. 2b and 4a, bottom). A previously reported crystal structure of a recombinant 8 fragment (PDB code 5JIE, https://doi.org/10.2210/pdb5JIE/pdb; residues Pro421-Ala483 in a-8 fusion protein) fits well into the distal fiber density (*5*) (Fig. 4a, bottom).

To address potential heterogeneity in fiber occupancy and flexibility, we performed additional 3D reconstructions using an expanded OrV dataset (Supplementary Table 1), including C1 refinement, relaxation of icosahedral symmetry, and focused refinement of the fivefold region (Fig. 4b and Supplementary Fig. 2b). Focused refinement improved local density, revealing a 28 residues segment corresponding to the linker region between the capsid a subunit and the 8 fiber (Fig. 4b-d). This linker forms a pentameric coiled coil. Within this region, we could assign several side chains, enabling sequence registration (Fig. 4c). The a-8 fusion protein, generated via a –1 ribosomal frameshift, has previously been estimated to be incorporated as one pentameric fiber per capsid based on its abundance in purified virions (Fig. 4d) (*21*). In our reconstructions, density corresponding to the fiber (a part of it), extending ∼80 Å, is consistently observed in both symmetry-imposed and C1 maps. Given that the occupancy of one fiber per capsid would be expected to be averaged out under icosahedral symmetry, the persistence of this signal across reconstructions suggests that the true incorporation level is likely higher than previously estimated. The linker region is highly conserved among several OrV-related nematode-infecting viruses that encode fiber-like structures, including Le Blanc and Santeuil viruses, suggesting conservation of a similar pentameric organization (Supplementary Fig. 11). Finally, AlphaFold3 modeling of five copies of full-length 8 protein (predicts an elongated, flexible assembly with a hinge region and Gly/Pro-rich linkers forming a tubular architecture, consistent with the relative flexibility and the incomplete density observed even in the focused-refined maps (Fig. 4e).

### Proposed assembly model of OrV

The structural proteins a, 8, and the a-8 fusion protein are encoded by RNA2, and all three can be incorporated into the capsid during virion assembly *in vivo*. However, recombinant expression of the capsid protein in *E. coli* produces OrV-VLPs together with a substantial proportion of soluble a dimers, which have been proposed to represent the only stable capsomere intermediate during capsid assembly (*2*).

Given the availability of both a and a-8 proteins for OrV capsid assembly *in vivo*, we used PISA software (*22*) to estimate the solvation free-energy gain upon interface formation (Δ*^i^G*) for penton assembly, together with the associated *P* value, which reflects interface specificity (*P* > 0.5, interface likely artifactual; *P* < 0.5, interface likely specific). As expected, fiber-containing pentons were more stable (Δ*^i^G* ≈ –34.7 kcal mol-^1^, *P* ≈ 0.35) than pentons lacking fibers (Δ*^i^G* ≈ –7.7 kcal mol-^1^, *P* ≈ 0.46), supporting a dual structural and functional role for the fiber. Structurally, we envisage that the a-8 fusion proteins promote penton assemblies facilitated by the self-oligomerization of the coiled-coil region of 8 (Fig. 5, stage 1). Functionally, incorporation of the fiber is also associated with viral infectivity (*5*). During morphogenesis, we propose that, in addition to a-8 pentamers, metastable pentamers composed of a subunits and/or mixed a/a-8 assemblies are stabilized by RNA binding, as suggested by the prominent genomic density underlying the penton (Fig. 3b). We cannot exclude the possibility that 8 also oligomerizes with the 8 domain of the a-8 fusion protein. The weak density observed for the linker region may reflect such an interaction, although it could also arise from the intrinsic flexibility of the linker. In this model, the flexible N-terminal regions of a and the a-8 fusion protein engage extensively with the RNA genome while retaining the conformational plasticity required to support capsid assembly (Fig. 5, stage 2). RNA segments could then interlace between neighboring penton assemblies promoting the encapsidation of the available RNA segments (Fig. 5, stages 3 and 4). Concomitantly, dimeric a proteins may associate with capsid-free regions of the assembling particle (Fig. 5, stage 5), bridging incomplete lattice patches through coordinated interactions between neighboring S and P domains. In this framework, the S domain is responsible for non-specific RNA segment encapsidation, whereas the P domains would progressively fine-tune intersubunit contacts and impose icosahedral lattice symmetry, thereby stabilizing and locking the capsid architecture into place (Fig. 5, stage 6).

**Figure 5.**
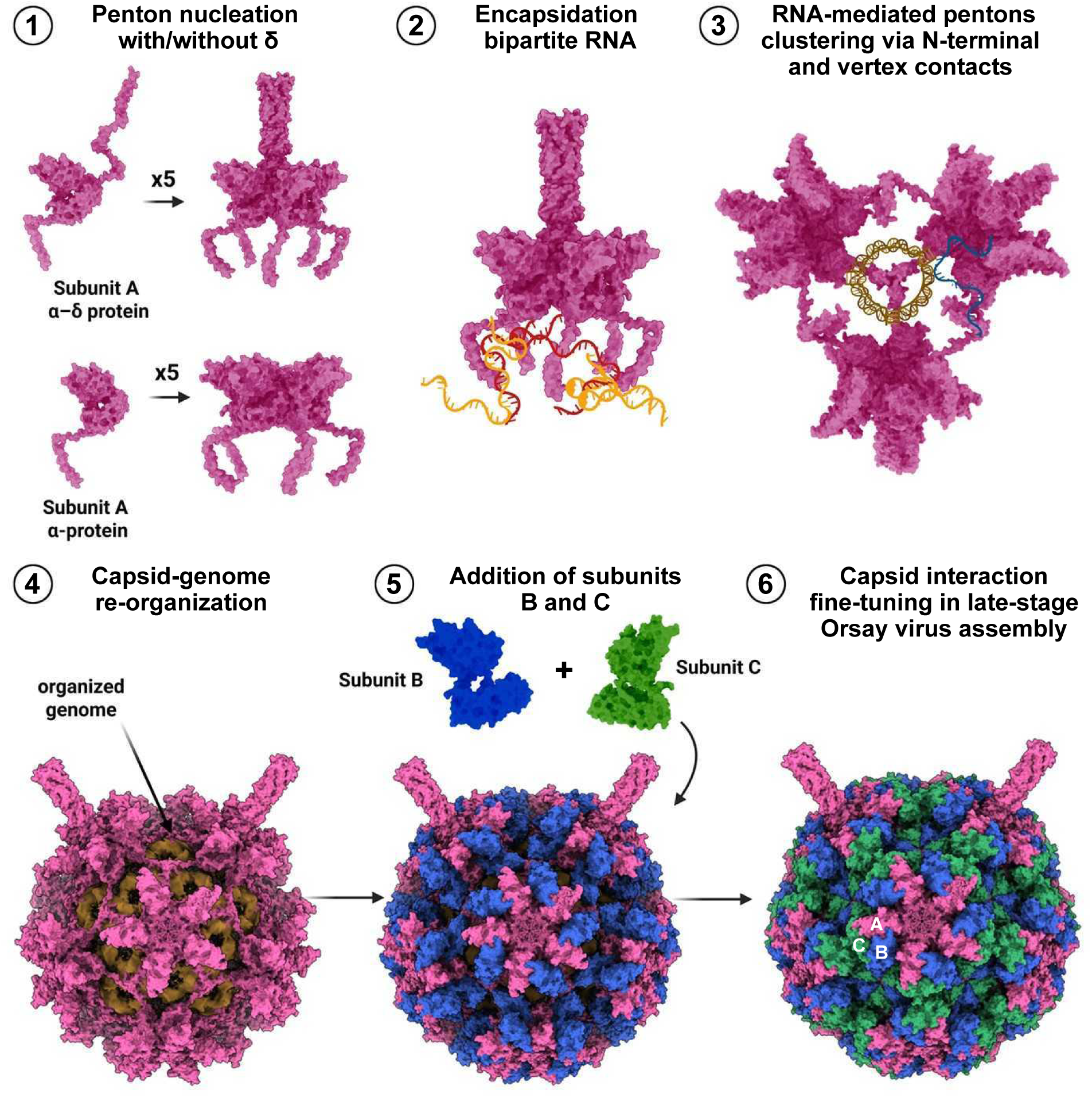
| Proposed steps model for OrV capsid assembly and genome encapsidation. (1) Penton nucleation by a or a-8 capsid proteins. (2) Encapsidation of bipartite RNA through electrostatic interactions with capsid N-terminal regions. (3) RNA-mediated clustering of pentons via N-terminal and vertex contacts. (4) Reorganization of the encapsidated genome beneath the capsid shell. (5) Addition of quasi-equivalent B and C capsid subunits during shell assembly. (6) Late-stage maturation involving fine-tuning of capsid interactions to generate the assembled virion. Capsid subunits A, B, and C are shown in pink, blue, and green, respectively; RNA is shown in orange/brown/navy-blue.

## DISCUSSION

Our study addresses how the bisegmented OrV infect cells and assembles virions despite unequal accumulation of its two genomic RNA segments, providing a framework to explore encapsidation strategies that extend beyond the canonical paradigm in which both genome segments are co-packaged into the same virion. Building on previous observations of asymmetric genome accumulation in the *C. elegans*-OrV pathosystem, we performed dose-response assays to assess the infectivity of individual virions.

Our results deviate from the expectation that each virion has a non-zero and dose-independent probability of initiating infection, indicating that productive replication requires coinfection by at least two virions. This finding suggests either the presence of a substantial fraction of defective particles or the existence of infectious particles that do not uniformly encapsidate a complete genome complement. To distinguish between these possibilities, we used cryo-EM analysis and determined the structure of purified native OrV. Reconstructions from both selected and expanded particle datasets reached near-atomic resolution, enabling detailed visualization of the quasi-equivalent A, B, and C capsid subunits that comprise the icosahedral asymmetric unit. Compared with the previously reported lower-resolution OrV VLP crystal structure (*2*), the native virion structure uncovers genome-capsid tethering regions, the fiber attachment site, and previously unresolved structural features. Notably, the Cys107-Cys195 disulfide bond observed in the B and C subunits is absent in the penton-forming A subunit, revealing a distinct structural state at the virion vertices that may facilitate both genome encapsidation and fiber accommodation. In contrast, subunits B and C, positioned near the threefold interfaces, retain Cys107-Cys195 disulfide bonds that stabilize the S_dom_. These subunits also exhibit disordered N-terminal regions in the resolved structure, likely interacting with the underlying genome but without following icosahedral symmetry, unlike the N-terminal arm of subunit A, which extends through adjacent B and C subunits and is resolved up to residue Ser30. Both density and secondary-structure predictions indicate that residues beyond Ser30 may adopt a-helical structure that by converging near the threefold axes form distinctive RNA anchoring sites. Overall, the capsid interior is predominantly electropositive, and *in vitro* nanoparticle reconstitution of the S_dom_ with RNA supports electrostatic interactions as a major determinant of genome encapsidation. We further assessed whether OrV formation yields heterogeneous populations with full, empty, or intermediate genome content, as reported for adeno-associated virus during ssDNA packaging (*23*). Two-dimensional classification of extracted virions and genome-focused classification of reconstruction particles did not identify any classes that could be confidently assigned as empty or partially filled virions. Most particles displayed comparable internal densities, while only a small subset (∼8%) exhibited distinct contrast within the genome region. The strongest density corresponds to a first layer of RNA closely apposed to the inner capsid surface. Given the available internal volume, the capsid could theoretically accommodate three to four genomic segments. However, although the present data do not resolve whether individual virions contain RNA1, RNA2, or mixed RNA1-RNA2 combinations, they argue against the existence of large populations of particles containing substantially different genome loads. Functionally, the requirement for two virions to achieve productive infection suggests that individual particles may not consistently encapsidate a full complement of RNA1 and RNA2 (with RNA2 ∼24% smaller in size than RNA1). If productive infection indeed requires the presence of at least one copy of both RNA1 and RNA2 within the same cell, stochastic genome assortment provides a simple quantitative explanation for the departure from the IAH observed in our dose-response experiments. If productive infection requires the presence of at least one copy of both RNA1 and RNA2, and encapsidation occurs without strict segment-specific selection, then the expected fraction of infectious particles can be estimated from simple combinatorial considerations. For a particle containing two genomic segments, random assortment predicts four equiprobable outcomes, of which only two contain both segment types, yielding a maximum infective fraction of 50%. For particles containing three segments, six out of eight possible combinations contain at least one copy of each segment, increasing the expected infective fraction to 75%. Therefore, for particles carrying on average two to three RNA molecules, as suggested by the cryo-EM volume estimates, stochastic encapsidation is expected to generate a substantial proportion of non-productive particles even in the complete absence of empty virions. Importantly, this estimate assumes equal intracellular abundances of RNA1 and RNA2. Any imbalance in segment availability, consistent with the preferential accumulation of RNA1 observed at early stages of infection, would further reduce the fraction of particles carrying a complete functional genome complement. Thus, the structural evidence for largely non-specific RNA encapsidation, together with the observed asymmetry in segment accumulation, provides a quantitative framework that is fully compatible with the cooperative infectivity inferred from the dose-response analyses.

These observations raise the question of whether the bisegmented OrV genome is randomly assorted or selectively packaged. Segmented RNA viruses exhibit diverse encapsidation strategies. In the *Comoviridae* subfamily (*e.g*., cowpea mosaic virus), RNA1 and RNA2 (RNA2 about 43% smaller in size than RNA1) are packaged into separate particles (*24*). In contrast, *Nodaviridae* (*e.g*., Flock House virus) can form non-covalent RNA heterodimers prior to packaging (*25*). More complex segmented viruses such as reoviruses and cystoviruses rely on highly selective packaging mediated by *cis*-acting RNA signals and protein-RNA interactions (*26*). Although our data do not directly address segment selectivity, they support a model in which OrV encapsidation is driven largely by nonspecific electrostatic interactions between RNA and the electropositive capsid interior. Consistent with this view, particle assembly occurs in the presence of RNA1, RNA2, or heterologous RNAs, arguing against strict recognition of segment-specific packaging signals. The semi-ordered genome organization and absence of large populations of empty or partially filled particles further support this model. The functional asymmetry observed during infection may therefore arise not from selective packaging itself, but from differences in intracellular replication dynamics and relative segment abundance. Greater availability of RNA1, for example, could bias encapsidation toward particles competent for replication initiation. In addition, compartmentalization of genome replication may further influence packaging by restricting RNA accessibility and limiting incorporation of non-viral or incomplete RNA molecules. Consistent with this possibility, Flock House virus remodels the outer mitochondrial membrane to generate spherular replication compartments that spatially organize RNA synthesis (*27*). Finally, we resolved the 8 fiber emerging from the fivefold vertices as a structural element of the a-8 fusion protein. Beyond its established role in host-cell attachment and infectivity (*5*), the 8 fiber linked to the a capsid protein may favour assembly toward 8-containing pentamers over a-only pentamers, while also promoting mixed a/a-8 assemblies. In the presence of RNA, these interactions likely contribute to a coordinated assembly pathway of penton-RNA complexes, further stabilized by capsid-RNA contacts, culminating in maturation of the particle through fine-tuning of capsid protein interactions.

Together, our findings support a model in which genome assortment emerges from the interplay between capsid assembly and RNA availability, with the latter likely shaped by segment-specific replication dynamics and intracellular organization. More broadly, this work provides a framework for understanding genome packaging in segmented RNA viruses that rely on stochastic rather than deterministic encapsidation.

## MATERIALS AND METHODS

### *C. elegans* strain maintenance

All *C. elegans* strains were cultured at 20 °C on standard Nematode Growth Medium (NGM) plates seeded with *E. coli* OP50. Strains utilized in this study included: JU2624 (*[myo-2::mCherry::unc-54; lys-3p::eGFP::tbb-2] IV*), derived from JU1580, for viral amplification and infection assays. ERT54 (*jyIs8 [pals-5p::GFP; myo-2p::mCherry] X*) for infection assays. SFE2 (*drh-1(ok3495) IV; mjIs228 [myo-2::mCherry::unc-54; lys-3p::eGFP::tbb-2] ?*), used exclusively for large-scale OrV purification.

### OrV stock preparation and quantification

Synchronized L1 JU2624 animals were inoculated on NGM plates with 50 µL of OrV JUv1580_vlc stock and incubated at 20 °C for five days. Animals were washed into M9 buffer (0.22 M KH_2_PO_4_, 0.42 M Na_2_HPO_4_, 0.85 M NaCl, 1 mM MgSO_4_), allowed to settle 15 min at room temperature, vortexed and pelleted (400 g, 2 min). The supernatant was clarified by two successive centrifugations at 21,000 g for 5 min, filtered through a 0.2 µm membrane, and stored at –80 °C. Viral RNA was extracted from 200 µL of clarified supernatant using the NZYtech Viral RNA Isolation Kit (Cat. No. MB40701; NZYtech, Lisboa, Portugal). RNA yield and purity were assessed by NanoDrop^TM^ Lite Plus (Cat. No NDLPLUSGL; Thermo Fisher Scientific; Waltham MA, USA). Genome copy number was determined by one-step RT-qPCR using the QIAGEN OneStep RT-PCR Kit (Cat. No. 210212; QIAGEN, Hilden, Germany) against a T7-transcribed RNA2 standard curve. RT-qPCR reactions (20 µL) contained 5 µL RNA template, 400 nM each primer, and 200 nM probe on an Applied Biosystems^TM^ QuantStudio 5 qPCR (Cat. No. 15791238; Thermo Fisher Scientific).

To generate an absolute quantification standard for RT-qPCR, we amplified a 1 kb fragment at the 3’ end of RNA2 from OrV JUv1580_vlc cDNA with Accuscript High-Fidelity Reverse Transcriptase (Cat. No. 200820; Agilent, Santa Clara CA, USA) using primers incorporating a T7 promoter sequence (Supplementary Table 2). The PCR product was gel-purified with MSB^®^ Spin PCRapace (Cat. No. 1020220300; Invitek Molecular GmbH, Berlin, Germany) and transcribed *in vitro* (T7 Polymerase; Cat No. RPOLT7-RO; Merck, Darmstadtm Germany). DNA was removed by Ambion^TM^ DNase I RNase-free (Thermo Fisher Scientific) treatment, and RNA integrity was confirmed on a denaturing agarose gel. RNA concentration was measured by NanoDrop (Thermo Fisher Scientific), and copy number was calculated using the EndMemo RNA Copy Number Calculator (https://www.endmemo.com/bio/dnacopynum.php). Serial 10-fold dilutions (10^2^-10^8^ copies/µL) of the transcript were prepared in nuclease-free water. These standards were included in each RT-qPCR run to convert *C_q_* values into absolute genome copy numbers.

### Synchronization, inoculation and smFISH

To obtain synchronized populations, NGM plates bearing gravid adults were washed gently with M9 buffer, removing larvae and adults while retaining eggs. Plates were incubated at 20 °C for 1 h to allow eggs to hatch; newly hatched L1 larvae were collected by washing with fresh M9. For infection, synchronized L1s (∼1,000 worms per plate for smFISH and ∼100 for infection scoring) were transferred to NGM plates seeded with OP50 and overlaid with 50 µL of OrV JUv1580_vlc at the indicated viral concentrations (infection scoring) or 2.8×10^9^ copies of RNA2 in the inoculum (smFISH). Plates were incubated at 20 °C for 8 or 24 h. smFISH was performed as previously reported (*11*).

### Infection scoring

Viral inocula were prepared as 10-fold serial dilutions in M9 to generate dose-response curves. Infection efficiency was assessed at 24 hpi by scoring GFP reporters (*pals-5p*::GFP in ERT54 or *lys-3p*::GFP in JU2624) under a Leica M165 FC stereomicroscope (Leica Microsystems, Wetzlar, Germany) equipped with a GFP filter set. Individual animals were scored as infected if anterior intestinal GFP signal was observed. Approximately 100 worms per condition were examined across five biological replicates. The percentage of infected animals (*I*) was calculated as *I* = (GFP-positive animals)/(total animals observed).

Dose-infection data were fitted to the IAH model using Levenberg-Marquard method of nonlinear regression using the R function nlsLM implemented in the minpack.lm package version 1.2-4. To evaluate strain (*S*) effects on the two model parameters, the following equation was fitted to the whole dataset: I(D) = 1 – exp[-(p + 8_p_S)D^(K+8KS)^], where *8ρ* and *8κ* quantify the strain-dependent effects on the parameters. The best fitting model was chosen according to the partial-*F* test method. Analyses were done in R version 4.5.0 under RStudio 2025.09.1 (Posit PBC, Boston MA, USA).

### OrV purification

Large-scale OrV was purified from infected SFE2 cultures following an adapted protocol (*4*). Briefly, *E. coli* OP50 were grown overnight, concentrated 10-fold in S medium, and inoculated with SFE2 worms (stable *lys-3p*::GFP reporter) at ∼1,000 individuals mL^−1^. Cultures were incubated 6 days at 20 °C, 160 rpm, with an additional viral boost on day 5. Infection was confirmed by GFP fluorescence, then cultures were chilled on ice for 30 min to settle worms and debris. Supernatants were clarified by centrifugation (30,000 g, 30 min, 4 °C), treated with 1% Triton X-100 (1 h, RT, 170 rpm). Virions were pelleted through ultracentrifugation (142,000 g, 2 h, 4 °C), resuspended in low-salt buffer (20 mM Tris-HCl pH 7.8, 100 mM NaCl, 0.1% 2-mercaptoethanol, 1 mM EDTA), incubated overnight and pelleted through ultracentrifugation (284,000 g, 2 h, 4 °C). The final pellet was resuspended in low-salt buffer, layered onto a 25-47.5% iodixanol step gradient, and centrifuged at 247,000 g for 16 h at 4 °C with minimal acceleration/deceleration. Fractions were collected manually, and viral RNA content was quantified by RT-qPCR to identify peak fractions (Supplementary Fig. 1). Selected fractions were pooled and concentrated by ultracentrifugation at 284,000 g for 2 h at 4 °C. The pellet was resuspended, adjusted to a final NaCl concentration of 1 M, and dialyzed overnight against PBS at 4 °C using Spectra-Por^®^ Float-A-Lyzer^®^ G2 devices (Cat. No. Z726060, Merck), with three buffer exchanges. The dialyzed material was subsequently concentrated tenfold using a Vivaspin^®^ 2 concentrator (Cat. No. VS02V2, Sartorius, Göttingen, Germany) at 8,000 g and 4 °C. Viral titers were quantified by RT-qPCR, and samples were stored at –80 °C until use.

### OrV RNA production

Complementary DNA (cDNA) of OrV and TuMV were synthesized from either purified OrV JUv1580_vlc RNA or from the infectious clone p35STunos that contains the complete sequence of TuMV strain YC5 (AF530055.2; (*28*)), respectively, using SuperScript^TM^ III Reverse Transcriptase (Cat. No. 18080085; Thermo Fisher Scientific) and gene-specific reverse primers targeting the 3’ end of each segment. PCR amplification was performed with Phusion^TM^ High-Fidelity DNA Polymerase (Cat. No. F534L; Thermo Fisher Scientific) using forward primers appended with a T7 promoter sequence (Supplementary Table 2). *In vitro* transcription was carried out with T7 RNA polymerase (Merck), followed by DNase I (Thermo Fisher Scientific) digestion. Transcripts were purified by phenol-chloroform extraction and ethanol precipitation. Integrity and size were confirmed by denaturing agarose gel electrophoresis (RiboRuler High Range RNA Ladder; Cat. No. SM1821; Thermo Fisher Scientific), and concentrations were determined spectrophotometrically (NanoDrop). RNA stocks were aliquoted and stored at –80 °C until use.

### Cryo-EM sample preparation and data collection

Purified OrV virions (0.1 mg mL^−1^ in 20 mM Tris-HCl pH 8.0, 300 mM NaCl) were applied (3 µL) to glow-discharged Quantifoil R1.2/1.3 Cu 300-mesh grids (Cat. No. N1-C14nCu30-01) and vitrified using a Leica EM GP2 system (Leica Microsystems) (14 °C, 95 % humidity, 40 s incubation, 1 s blot) by plunge-freezing into liquid ethane. Grid quality and particle distribution were initially examined on a JEM-2200FS microscope (JEOL Ltd.) equipped with a Gatan K2 detector (Gatan. Inc.). High-resolution cryo-EM data were collected at the Electron Bio-Imaging Centre (eBIC, Diamond Light Source) using a Titan Krios G4 transmission electron microscope (Thermo Fisher Scientific) operated at 300 kV and equipped with a Gatan K3 direct electron detector in counting mode. Automated acquisition was performed with EPU software (Thermo Fisher Scientific) at a nominal magnification of 105,000×, corresponding to a calibrated super-resolution pixel size of 0.829 Å. Two independent datasets comprising 26,492 and 22,858 movies were collected (Supplementary Table 1). Movies were recorded as 50-frame image stacks using fringe-free imaging with beam-tilt compensation and Aberration-Free Image Shift (AFIS) enabled. The total accumulated electron dose was 46.0 and 50.0 e– A-^2^ for datasets A and B, respectively, with a defocus range of –0.8 to –2.4 µm.

### Image processing and 3D reconstruction

For dataset A, movie-frame alignment and dose weighting were performed using MotionCor2, and contrast transfer function (CTF) parameters were estimated with CTFFIND4 (*29*,*30*). Particles were automatically picked with crYOLO, extracted in 640-pixel boxes, and fourfold binned prior to image processing (*31*). Two-dimensional and three-dimensional classifications in RELION-v4 identified a subset of 38,101 high-quality particles, which were refined under icosahedral symmetry to yield a reconstruction at 2.5 Å resolution based on the 0.143 gold-standrad Fourier shell correlation (FSC) criterion (Supplementary Fig. 2a and Supplementary Table 1) (*16*).

To investigate asymmetric features, datasets A and B were subsequently combined and processed in cryoSPARC v4.7 (*17*). Reconstructions were generated using three refinement strategies: icosahedral symmetry, fully asymmetric (C1) refinement, and symmetry-relaxed refinement, yielding maps at 2.6, 3.2, and 3.3 Å resolution, respectively (Supplementary Fig. 2b). Refinement around the fivefold vertices improved visualization of the 8-fiber stalks and RNA-capsid interaction sites. For genome classification and analysis, we performed reference-free 2D classification of particles from the combined datasets A and B and focused classification of internal densities from combined particles contributing to the OrV 3D icosahedral reconstruction. Capsid and fiber densities were subtracted before genome focused classification.

### Model building and refinement

The OrV IAU was manually fitted into the 2.5 Å cryo-EM map using COOT v.0.9.8.7, with the 3.2 Å crystallographic structure (PDB code 4NWV; https://doi.org/10.2210/pdb4NWV/pdb) as an initial reference model (*2, 32*). The capsid-only TAU model, comprising the three a proteins (chains A, B, and C), was refined against the corresponding boxed cryo-EM density using Phenix v1.20.1-4487 software (*33*). Refinement included rigid-body fitting, real-space minimization with secondary-structure restraints, and final atomic displacement parameter (ADP) refinement (initial isotropic B-factor of 40 Å^2^) (Supplementary Table 1). Water molecules were added iteratively during refinement using criteria that included a maximum distance of 3.4 Å from the atomic model and a density threshold greater than 1.5a (boxed map in COOT v.0.9.8.7). The final model was expanded by 60-fold NCS to generate the complete virion, after which the 60 subunits were independently rigid-body refined against the full cryo-EM density map.

Tail-fiber densities in the locally refined C1 map were interpreted by initially rigid-body fitting the previously reported structure (PDB code 5JIE; https://doi.org/10.2210/pdb5JIE/pdb) into the corresponding density using ChimeraX (*20*). The C-terminal linker of subunit A connecting the a and 8 proteins was subsequently built *de novo* from residues Tyr385 to Pro421 using a predicted AlphaFold3 model of the penton a-8 fragment as a template, guided by the presence of bulky side-chain densities for sequence registration (*18*). Peripentonal 8 fragments were then extracted to generate locally boxed cryo-EM densities, followed by geometry regularization through five macrocycles of highly constrained real-space refinement in Phenix (Supplementary Table 1).

The refined 8 fragment of chain A (residues 387-452) was merged with the capsomer model derived from the icosahedral reconstruction reported in this study (EMD-57282; PDB 29QM), and residues 386-388 were regularized in COOT. The resulting model and the corresponding C1 OrV density map, locally refined around the penton linker region, were deposited in the Protein Data Bank and Electron Microscopy Data Bank under accession codes PDB 32NR and EMD-59028, respectively.

### Expression and purification of the S-domain and its N-truncated mutant

Two truncated constructs of the OrV capsid protein were engineered: S_dom_ (residues 1-211), comprising the N-terminal arm and shell domain, and ΔN-S_dom_ (residues 42-211), containing only the shell domain. Both constructs were cloned into a modified pDB_His_Gst vector (DNASU repository, https://dnasu.org/DNASU/Home.do) in which the N-terminal His_6_-GST fusion tag had been removed. A C-terminal 5×His tag was introduced into each construct to facilitate purification. Proteins were expressed in *E. coli* BL21 (DE3) cells (Cat. No. CMC0014; Merck) grown in LB medium at 37 °C to an OD_600_ of ∼0.8, induced with 1 mM IPTG, and incubated for 16 h at 18 °C. Cell pellets were resuspended in lysis buffer containing 25 mM Tris-HCl (pH 7.5), 10% glycerol, and 40 mM imidazole, supplemented with either 300 mM NaCl for S_dom_ or 500 mM NaCl for ilN-S_dom_. Cells were lysed by sonication and clarified by centrifugation. Proteins were purified by Ni-NTA (Cat. No. 88222; Thermo Fisher Cientific) affinity chromatography followed by size-exclusion chromatography. S_dom_ was further purified on a Hi load 16/600 Superdex 200 (Cat. No. 28989335; Cytiva) column equilibrated in 25 mM Tris-HCl (pH 7.5) and 300 mM NaCl, whereas ilN-S_dom_ was purified on a HiLoad 16/600 Superdex 75 (Cat. No. 28989333; Cytiva) column equilibrated in 25 mM Tris-HCl (pH 7.5) and 500 mM NaCl.

### Oligomeric analysis of S_dom_ and ΔN-S_dom_ proteins

Oligomeric state was first assessed from the elution profile and SDS-PAGE of purified fractions for both S_dom_ and ΔN-S_dom_. Additional biophysical analyses were performed only for S_dom_, including NMR spectroscopy and chemical crosslinking (*34*). For ^1^H NMR, S_dom_ was examined at 10 µM and 130 µM dissolved in D_2_O, 25 mM Tris pH 7.5, 0.5 mM NaCl, either alone or in complex with RNA at a 10:1 molar ratio, using an 800 MHz AVANCE III spectrometer at 298 K (Bruker). For crosslinking, purified S_dom_ was incubated with 0.1% or 0.05% glutaraldehyde at 30 °C in buffer containing 20 mM Tricine (pH 8.3), 50 mM NaCl, 20% glycerol, 5 mM MgCl_2_, and 3 mM DTT, at final protein concentrations of 20, 10, 5, or 2 µM. Samples were preincubated on ice for 15 min before glutaraldehyde addition, and reactions were quenched after 30 min with 1 M Tris-HCl (pH 6.8). Crosslinked products were resolved by SDS-PAGE and visualized with Coomassie Brilliant Blue staining.

### RNA and DNA binding assays with S_dom_ and ΔN-S_dom_

Interaction assays were performed to evaluate the binding of constructs, S_dom_ and ΔN-S_dom_ with viral RNA fragments (RNA1 and RNA2). Purified RNA1 and RNA2 segments of OrV were used individually or in combination. For S_dom_, assays were conducted at two protein concentrations: 43.4 µM and 14.8 µM. Each protein preparation was incubated with RNA1 (0.1 µM or 0.2 µM), RNA2 (0.1 µM or 0.2 µM), or a mixture of RNA1 (0.1 µM) and RNA2 (0.1 µM) to yield a final combined RNA concentration of 0.2 µM. ΔN-S_dom_ was assayed at 14.8 µM protein with the same RNA concentrations and combinations. All materials, including buffers, were rendered RNA-free by autoclaving. The interaction buffer consisted of 25 mM Tris-HCl, 150 mM NaCl, and 1 mM CaCl_2_ (pH 7.5). Controls included protein-only and RNA-only samples. For experimental treatments, protein and RNA were mixed on ice, followed by incubation at 30 °C for 10 min. Samples were then returned to 4 °C until vitrification and particle size determination. Vitrification parameters were identical to those described previously for cryo-EM sample preparation. The grids were observed using a JEM-2200FS transmission electron microscope equipped with a Gatan K2 Summit detector at a nominal magnification of 30×.

The same protocol was applied to additional nucleic acids: (*i*) TuMV RNA (∼6 kb), (*ii*) short synthetic DNA oligonucleotides of 100, 50 and 39 nucleotides designed to adopt distinct secondary structures, and (*iii*) a 2 kb random-sequence DNA fragment.

### EMSA

Unlabeled RNA1 and RNA2 were analyzed individually or mixed together. Two types of preparations were prepared: (*i*) straight mix of RNA1 and RNA2 with or without incubation time (at 20 °C for 25 min); (*ii*) individual RNAs were heat-denatured at 80 °C for 2 min and allowed to refold before mix with or without incubation time (at 20 °C for 25 min). Binding reactions were prepared in 1× binding buffer containing 20 mM HEPES (pH 7.5), 30 mM KCl, 5 mM MgCl_2_, 5% (v/v) glycerol, 100 mM DTT, and RiboLock RNase Tnhibitor (1 U µL-^1^; Cat. No. 10859710, Thermo Fisher Scientific).

To assess binding of the OrV capsid protein FP construct to nucleic acids, EMSAs were also performed using synthetic DNA oligonucleotides of 100, 50, and 39 nucleotides designed to adopt distinct secondary structures. Following the binding reactions, samples were mixed with loading dye and resolved on 1% agarose gels under native conditions. RNA samples were separated in 1× TBE buffer and stained with SYBR Gold (Cat. No. S11494, Thermo Fisher Scientific), whereas DNA samples were resolved in 1× TAE buffer containing 1× SYBR Safe DNA stain (Cat. No. S33102, Thermo Fisher Scientific). Electrophoresis was performed at 80-100 V for 1 h at room temperature. Gels were visualized using an ImageQuant LAS 4000 system (GE Healthcare/Amersham). Shifts in nucleic acid mobility relative to nucleic-acid-only controls were interpreted as evidence of protein-nucleic acid complex formation.

## Acknowledgements

We thank the EM Platform at the CIC bioGUNE for infrastructural support and preliminary cryo-grid screening. We acknowledge Diamond Light Source (UK) for access and support of the cryo-EM facilities at eBIC [under proposal BI31586], funded by the Wellcome Trust, MRC and BBRSC. We thank Francisca de la Iglesia (I2SysBio) for excellent technical assistance and Ron Geller and Cristina Fernández (I2SysBio) for help with ultracentrifugation.

## Data Availability

The cryo-EM maps generated in this study have been deposited in the Electron Microscopy Data Bank under accession codes EMD-57282 (icosahedral reconstruction) and EMD-59028 (C1 reconstruction locally refined). The corresponding atomic coordinates have been deposited in the Protein Data Bank under accession codes 29QM and 32NR, respectively.

## Supplementary Materials

Supplementary material include:

Supplementary Figures 1 to 11

Supplementary Tables 1 to 2

