## Supplemental Material for "Native virion structure reveals stochastic genome assortment underlying dose-dependent Orsay virus infection"

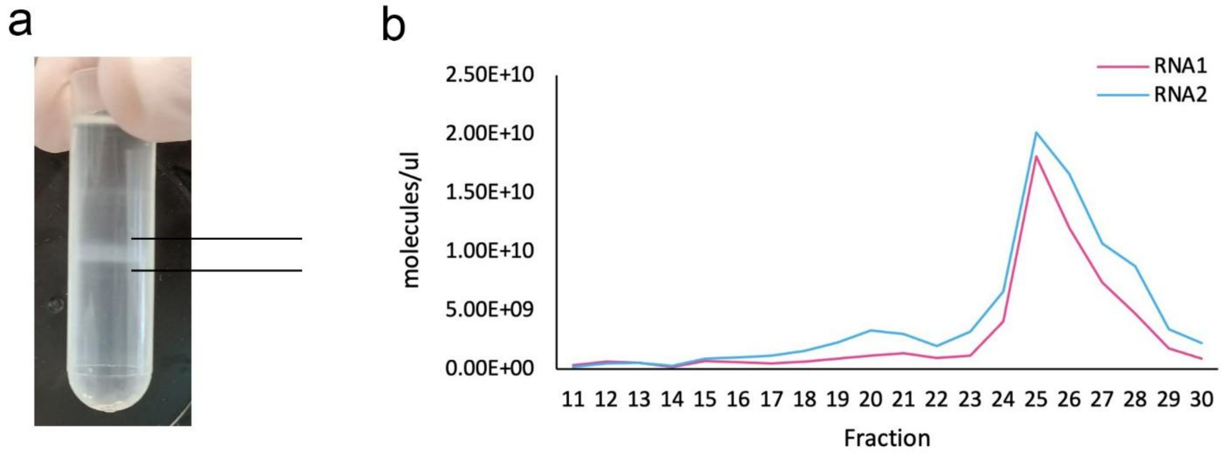


**Supplementary Figure 1 | OrV purification and RNA segment analysis.** (**a**) Representative density gradient after ultracentrifugation showing a visible translucent band (black lines) corresponding to particle-containing fractions. (**b**) Distribution of the two viral genomic RNA segments (RNA1, magenta; RNA2, blue) across gradient fractions, quantified as RNA molecules per microliter. Both RNA segments co-sediment, with peak abundance detected in fractions 25–26, corresponding to the lower particle band observed in the gradient. RNA copy numbers were determined by quantitative RT-PCR.

a

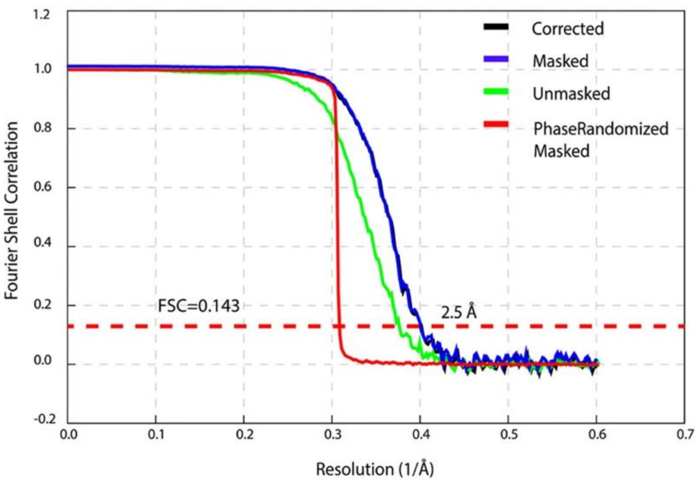

Corrected
Masked
Unmasked
PhaseRandomized Masked
Fourier Shell Correlation
Resolution (1/Å)
FSC=0.143
2.5 Å


b

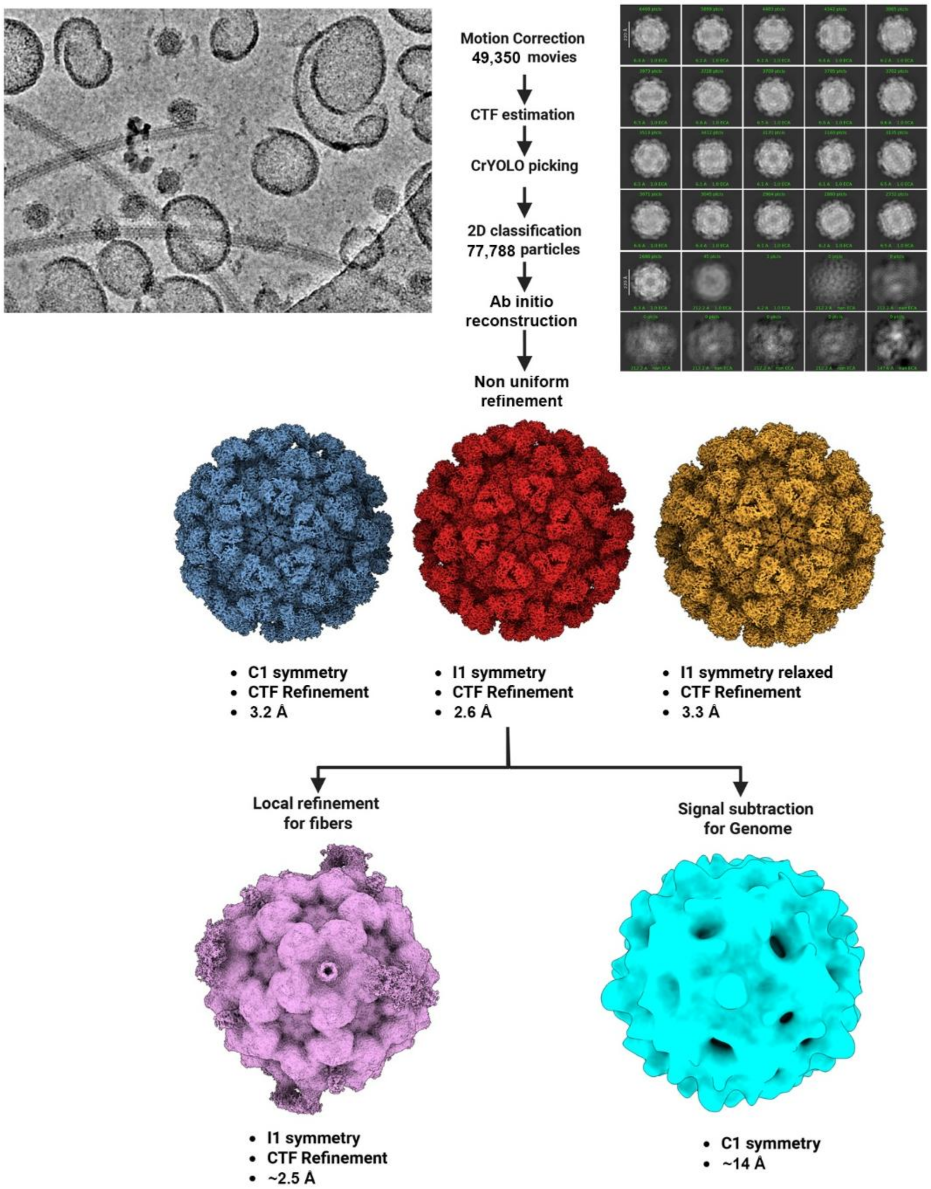

Motion Correction 49,350 movies
CTF estimation
CrYOLO picking
2D classification 77,788 particles
Ab initio reconstruction
Non uniform refinement
C1 symmetry
CTF Refinement
3.2 Å
I1 symmetry
CTF Refinement
2.6 Å
I1 symmetry relaxed
CTF Refinement
3.3 Å
Local refinement for fibers
Signal subtraction for Genome
I1 symmetry
CTF Refinement
~2.5 Å
C1 symmetry
~14 Å

**Supplementary Figure 2 | OrV resolution and image processing workflow under different symmetry conditions.** (**a**) Gold-standard Fourier shell correlation (FSC) curves for the final icosahedral reconstruction with 38,101 contributing particles, showing masked (blue), unmasked (green), corrected (black), and phase-randomized masked (red) estimates. The 0.143 criterion (dashed line) indicates an overall resolution of ~2.5 Å. (**b**) Overview of the cryo-EM data processing pipeline for the larger set of particles and 3D reconstructions under different symmetry and processing conditions. Representative micrograph of OrV particles (top left) and workflow from motion correction, CTF estimation, particle picking (top centre), and 2D classification (top right) to ab initio reconstruction and non-uniform refinement (center and bottom).

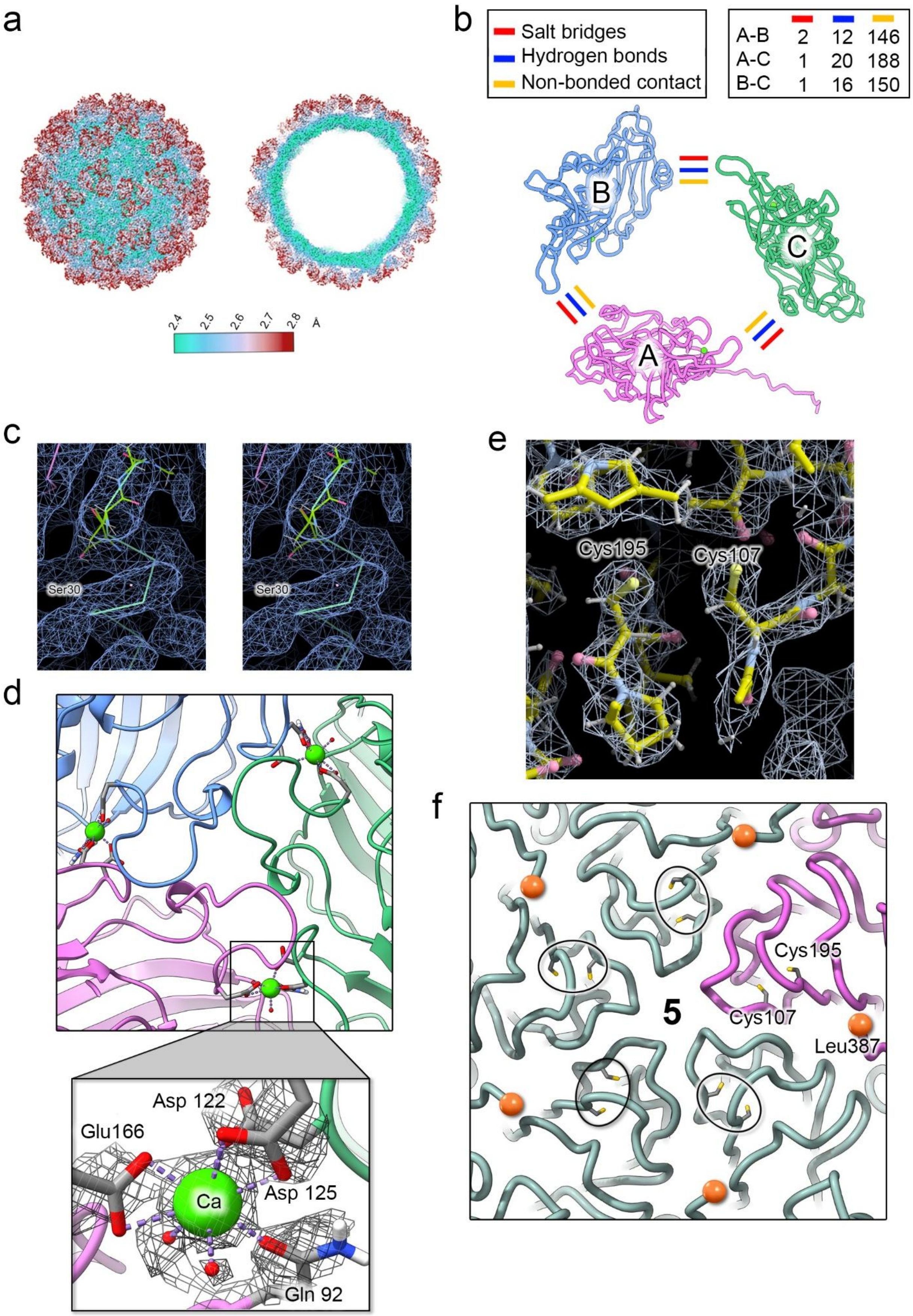

a
2.4
2.5
2.6
2.7
2.8
Å
b
Salt bridges
Hydrogen bonds
Non-bonded contact
A-B 2 12 146
A-C 1 20 188
B-C 1 16 150
B
C
A
c
Ser30
Ser30
d
Asp 122
Glu166
Ca
Asp 125
Gln 92
e
Cys195
Cys107
f
Cys195
5
Cys107
Leu387

**Supplementary Figure 3 | Inter-subunit interactions, structural features of the capsid and metal coordination.** (**a**) Surface (left) and cross-sectional (right) views of the cryo-EM map, colored by local resolution (legend below), illustrating higher resolution in the capsid S domain and lower resolution in the more flexible P domain. (**b**) Schematic of the trimeric assembly showing the three subunits (A, B, and C) and their interface interactions; the green dot visible within the structure marks the $Ca^{2+}$ ion. Salt bridges (red), hydrogen bonds (blue), and non-bonded contacts (yellow) are indicated. The accompanying table summarizes the number of interactions observed at each interface (A–B, A–C, and B–C). (**c**) Stereoview of the density at the N-terminus of subunit A in COOT. The cryo-EM map was low-pass filtered to 4 Å and is shown as a blue mesh (~1.3σ). Residues up to Ser30 from the deposited model are shown as sticks, whereas the cyan backbone represents additional residues fitted into the density, forming a putative α-helix. (**d**) Close-up of inter-subunit $Ca^{2+}$-binding sites. Inset shows the coordination environment of a representative $Ca^{2+}$ ion involving Glu166, Asp122, Asp125, Gln92, and two water molecules (red spheres); density is shown as a grey mesh. (**e**) Cryo-EM density (~1.8σ) showing the absence of a disulfide bond between Cys107 and Cys195, with chain A fitted into the density. (**f**) Structural organization around the fivefold axis. Cys107 and Cys195 precede the loops forming the fivefold plateau, whereas Leu387 is the final C-terminal residue (orange spheres) before the flexible linker of the δ protein.

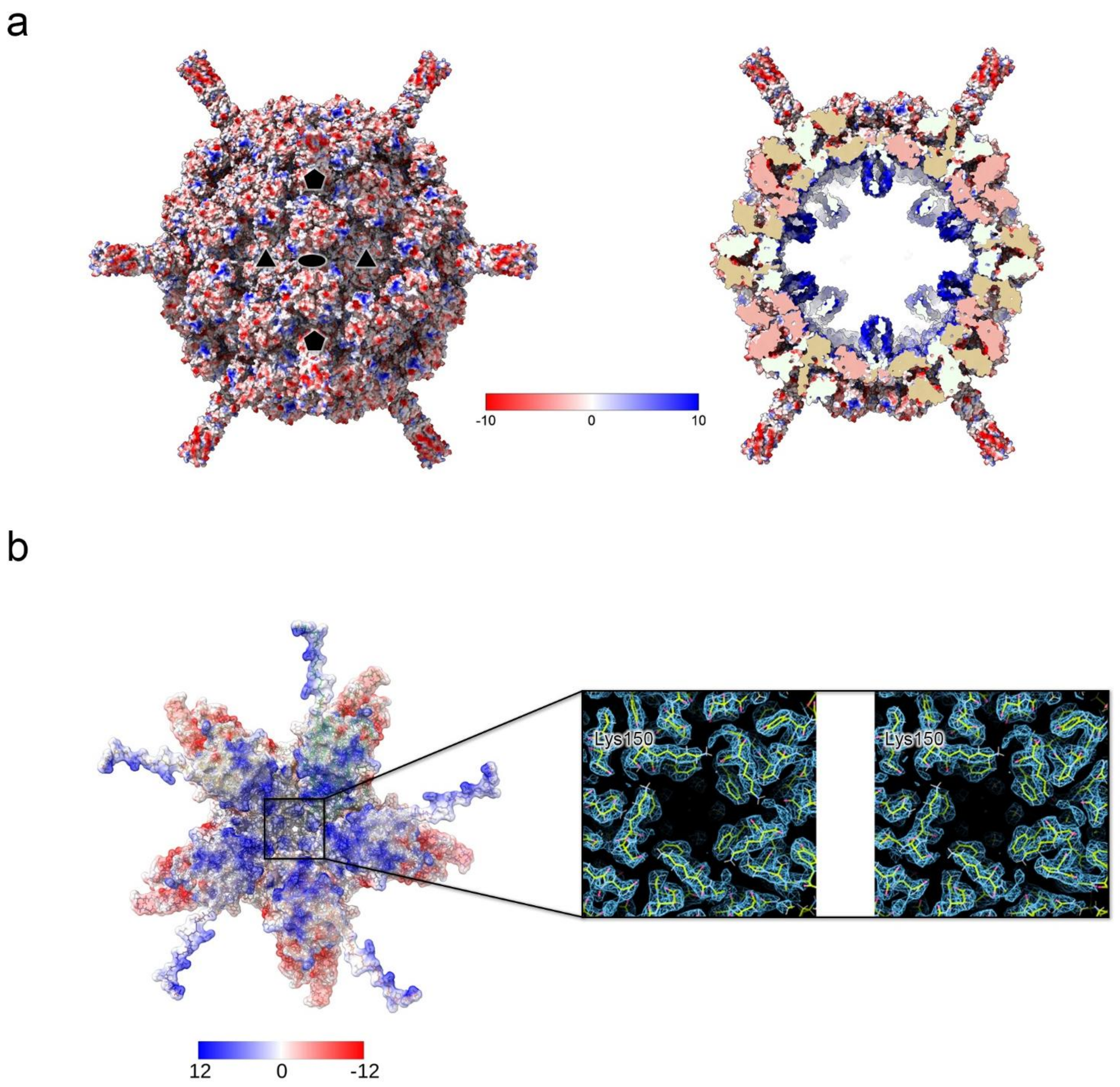


**Supplementary Figure 4 | Electrostatic rendering of the entire OrV and the region beneath the vertex.** (**a**) Surface electrostatic representation of the viral particle calculated and displayed in ChimeraX. Left: external view of the capsid surface colored according to electrostatic potential (scale from −10 to +10, red to blue). Right: cross-sectional view showing the internal organization, with positively charged regions lining the inner capsid surface and putative RNA–protein interaction interfaces highlighted. Five-, three-, and two-fold symmetry axes are indicated by two black pentagons, two black triangles, and one black oval, respectively. (**b**) Isopotential surface highlighting the positively charged region beneath the five-fold axis that anchors the viral genome beneath the vertex, giving rise to the bell-shaped averaged density. The black square marks the region enlarged in the inset, which shows a stereoview of the atomic model in stick representation with atoms colored by element.

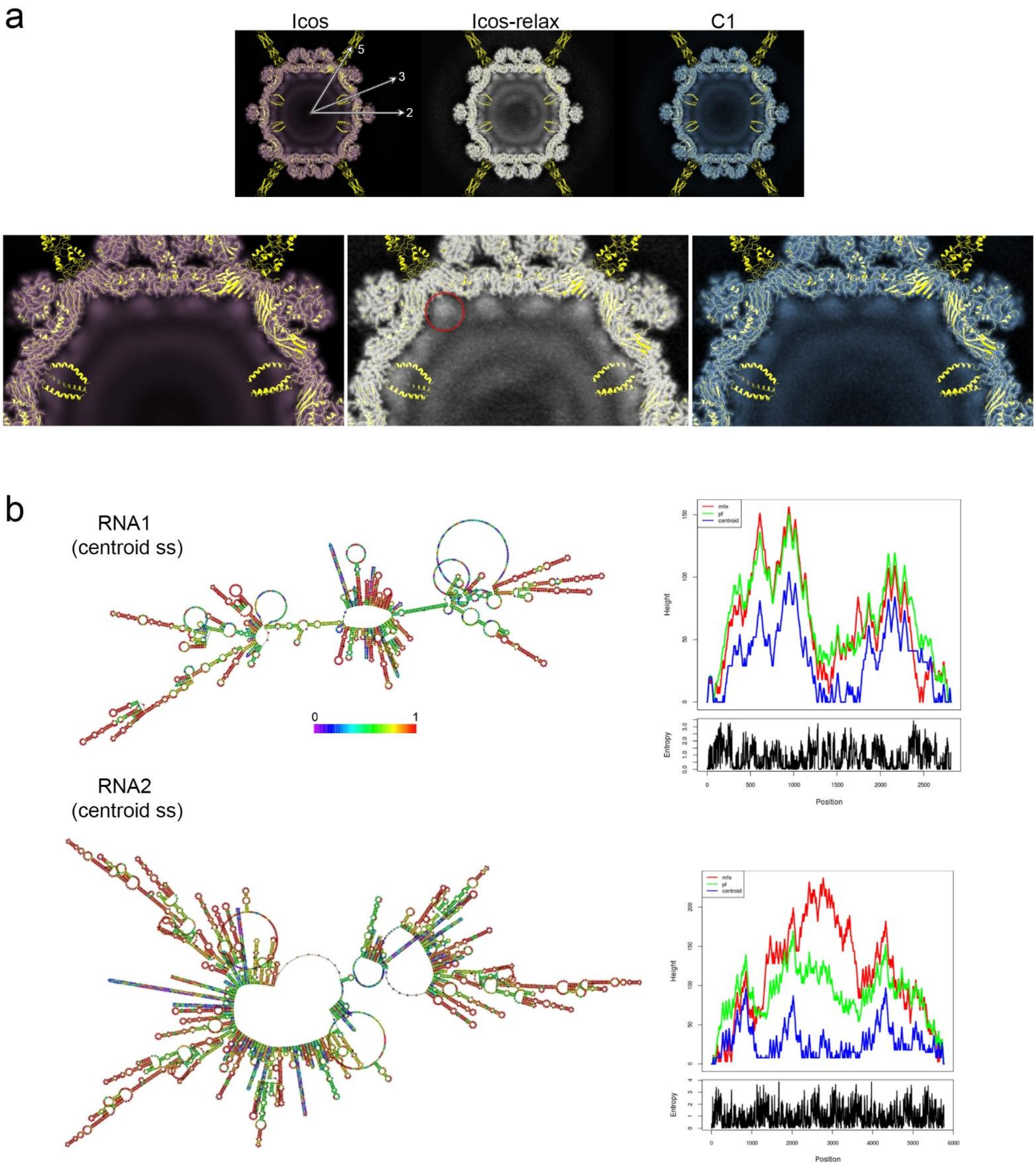


**Supplementary Figure 5 | Averaged OrV genome organization and RNA secondary-structure predictions.** (**a**) Top, unsharpened reconstructions visualized as central projections in ChimeraX obtained using icosahedral symmetry (Icos), icosahedral symmetry and then with relaxed symmetry (Icos-relax), and no symmetry imposed (C1) with superimposed the capsid atomic model; the N-terminal helices extending inside the capsid have been predicted by Alphafold3. Enlarged views (bottom) of the top half sections show illustrating that the internal genome density is relatively similar across all three reconstructions despite the different refinement strategies. The red circle highlights the bell-shaped density beneath the fivefold vertices (and across all reconstructions). (**b**) Secondary structure predictions and ensemble properties of genomic RNA segments using RNAFold software. Left: predicted centroid secondary structures of RNA1 (top) and RNA2 (bottom). Nucleotides are colored according to base-pairing probabilities (scale from 0 to 1), where warmer colors indicate higher pairing confidence and cooler colors indicate lower confidence or structural flexibility. Right: corresponding mountain plots (upper panels) showing structural profiles along the sequence for the minimum free energy (MFE, red), thermodynamic ensemble (partition function, green), and centroid (blue) structures. The *x*-axis represents the nucleotide position, and the *y*-axis indicates the number of base pairs enclosing each nucleotide ("mountain height"). Agreement between curves indicates well-defined structural regions, whereas divergence reflects structural heterogeneity within the ensemble. Lower panels: positional entropy profiles derived from base-pairing probabilities, providing a measure of structural variability at each nucleotide position. Higher entropy values indicate increased conformational flexibility and the presence of alternative base-pairing interactions.

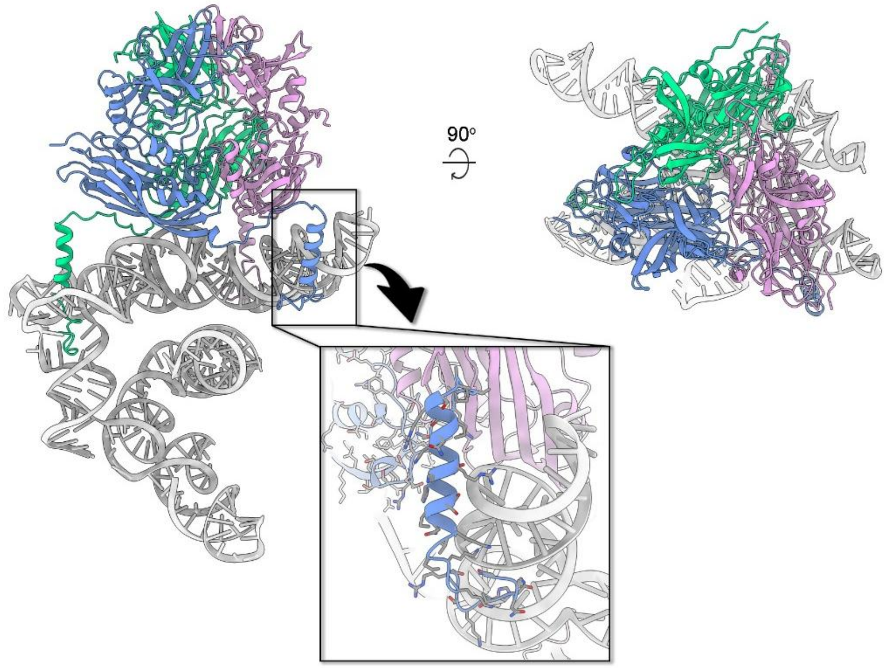


**Supplementary Figure 6 | Hypothetical Alphafold3-predicted model of capsid−RNA interactions beneath the threefold.** The capsomer is shown in cartoon and colored in blue, green, and light-magenta (as inset in Fig. 2c), while nucleic acids are depicted in grey. The left panel illustrates the complex bound to a nucleic acid scaffold, highlighting an extended interface between the protein surface and RNA architecture. The right panel shows the same assembly rotated by 90°, emphasizing the spatial organization of subunits around the nucleic acid. The boxed region is enlarged below and reveals a putative interaction site in which an α-helical segment inserts into the RNA groove, consistent with a shape- and charge-complementary mode of capsid–RNA recognition.

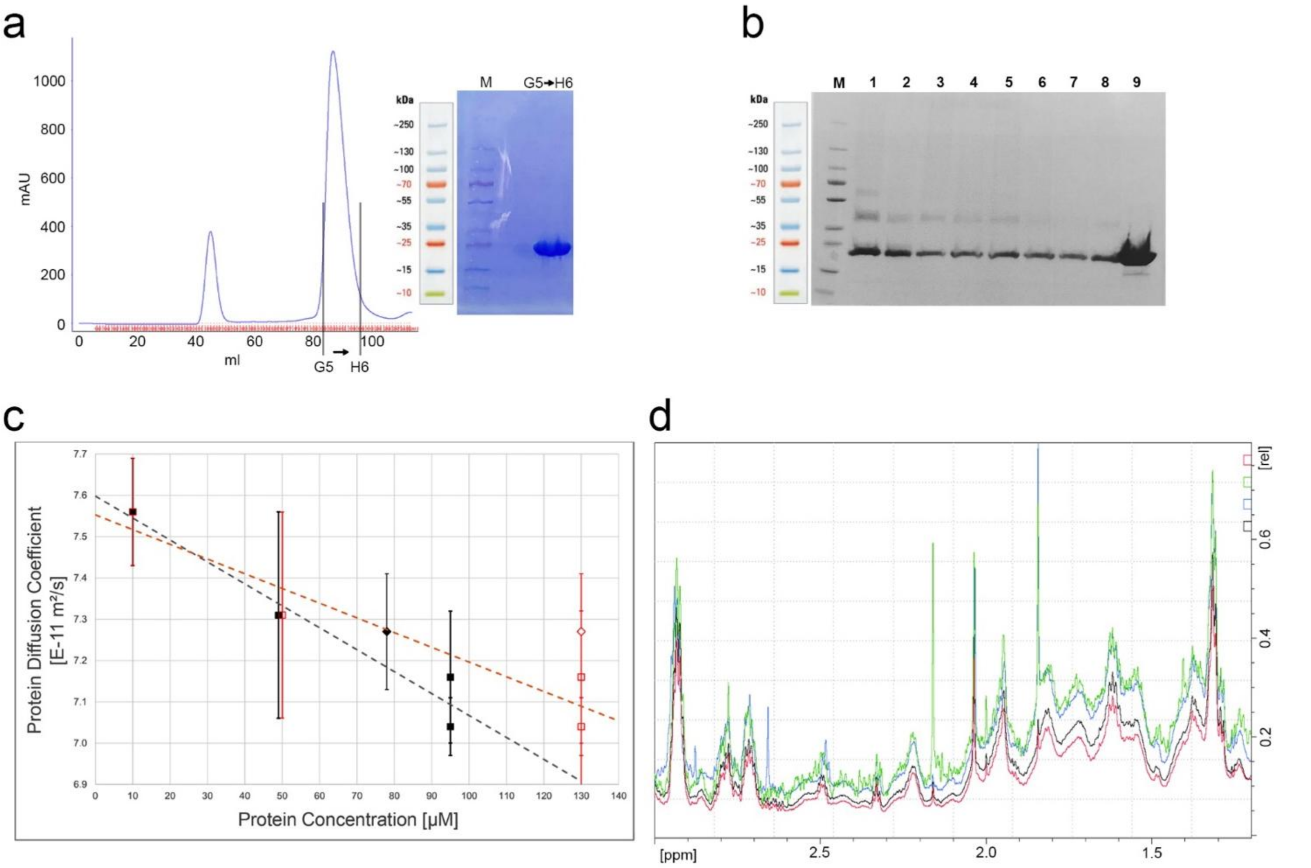


**Supplementary Figure 7 | S-domain biochemical and biophysical characterization and oligonucleotide interaction.** (**a**) Size-exclusion chromatography profile of purified S domain ($S_{dom}$), showing two principal elution peaks (fractions G5 and H6). Inset, SDS–PAGE analysis of pooled G5+H6 fractions confirming protein purity. (**b**) Cross-linking SDS–PAGE analysis of fractions with different concentration of $S_{dom}$ and percentages of glutaraldehyde for crosslinking. Lane M, molecular weight marker; lanes 1–9 correspond to: (1) 20 µM $S_{dom}$ with 0.1% glutaraldehyde; (2) 20 µM $S_{dom}$ with 0.05% glutaraldehyde; (3) 10 µM $S_{dom}$ with 0.1% glutaraldehyde; (4) 10 µM $S_{dom}$ with 0.05% glutaraldehyde; (5) 5 µM $S_{dom}$ with 0.1% glutaraldehyde; (6) 5 µM $S_{dom}$ with 0.05% glutaraldehyde; (7) 2 µM $S_{dom}$ with 0.1% glutaraldehyde; (8) 2 µM $S_{dom}$ with 0.05% glutaraldehyde; and (9) 20 µM $S_{dom}$ control (no glutaraldehyde). (**c**) Concentration-dependent diffusion coefficients (D) of $S_{dom}$ measured by $^{1}$H NMR at 298 K (800 MHz). Open squares represent nominal protein concentrations (10, 50, and 130 µM), whereas filled squares indicate effective NMR-observable concentrations after correcting for signal loss due to self-association. Error bars denote s.d. of 14 independent protein resonances. Linear fits (dashed lines) show increasing diffusion coefficients with increasing concentration, consistent with reversible self-association and depletion of slowly diffusing, NMR-invisible aggregates (> 102 kDa). Diamonds indicate measurements after addition of RNA (6.8 kb; $S_{dom}$:RNA ≈ 10:1), which caused an additional ~16% reduction in NMR-observable protein and an increased average diffusion coefficient, consistent with further sequestration of $S_{dom}$ into large RNA-associated assemblies. (**d**) $^{1}$H NMR spectra of $S_{dom}$ (3.3–1.2 ppm) acquired at 10 µM (green), 50 µM (blue), and 130 µM (black), scaled to compensate for dilution. The ~27% signal loss at 130 µM indicates formation of NMR-invisible aggregates (> 102 kDa), corresponding to an effective observable concentration of ~95 µM. Addition of RNA (red; 6.8 kb, $S_{dom}$:RNA ≈ 10:1) resulted in a further ~16% decrease in signal intensity, indicating enhanced $S_{dom}$ aggregation upon RNA binding.

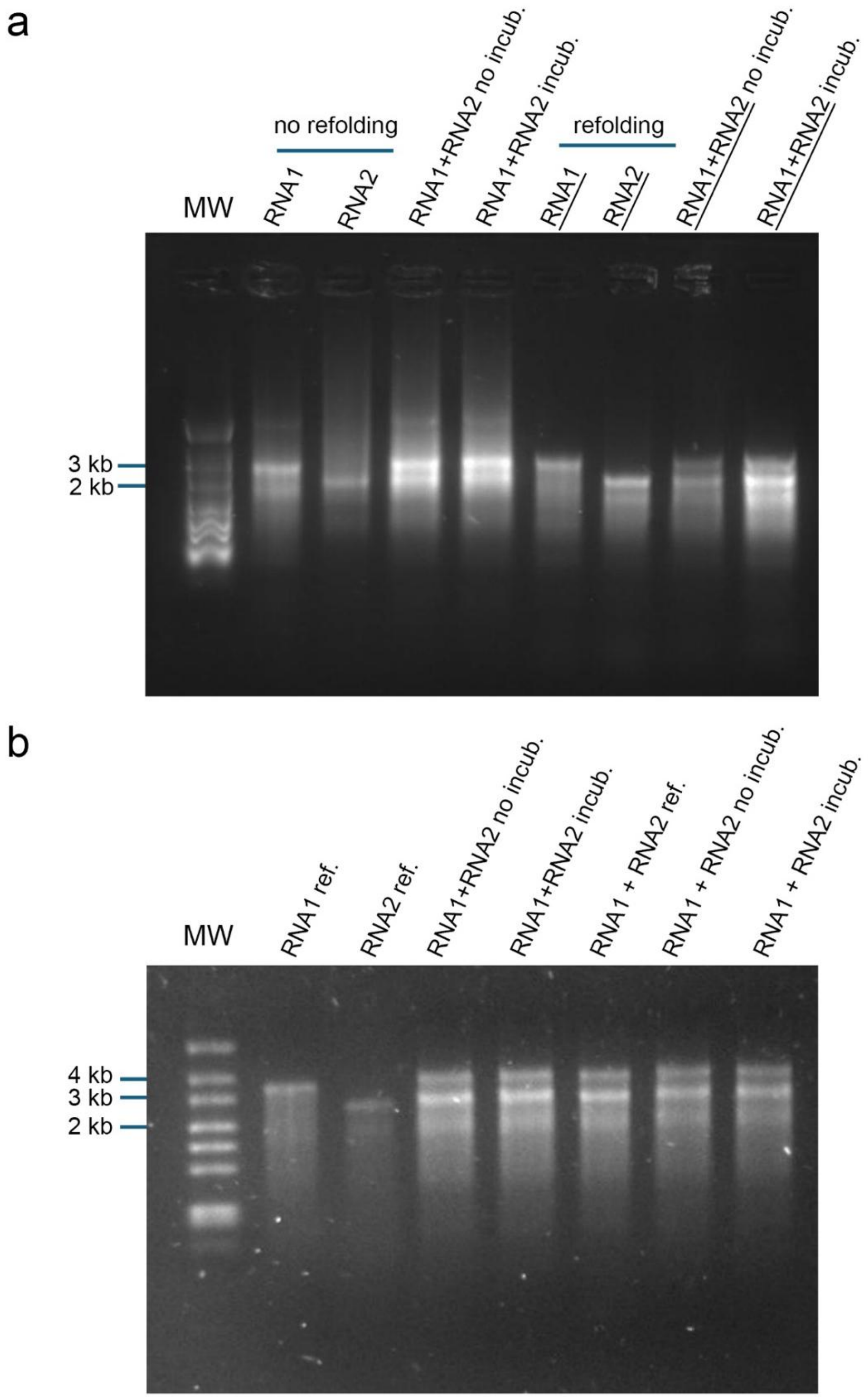


**Supplementary Figure 8 | Native gel analysis of RNA1 and RNA2 segments.** (**a**) Native agarose gel electrophoresis of RNA1, RNA2, and their equimolar mixture under non-refolded and refolded conditions (MW, molecular weight marker). RNA1 and RNA2 were analyzed individually either without refolding or after refolding. RNA mixtures were analyzed immediately after mixing (no incubation) or following incubation to assess RNA complex formation. (**b**) Native agarose gel electrophoresis performed as in (a). The first four lanes (excluding the molecular weight marker) replicate individually refolded RNA1 and

RNA2 mixed immediately before loading or after incubation as in (a). In the last three lanes, RNA1 and RNA2 were mixed prior to refolding and analyzed either immediately or after incubation. Comparable electrophoretic mobility of the RNA mixtures under all conditions, together with the absence of slower-migrating species, indicates that RNA1 and RNA2 do not form higher-order RNA complexes under the conditions tested. The positions of the 2 kb, 3 kb and 4 kb molecular weight markers are indicated.

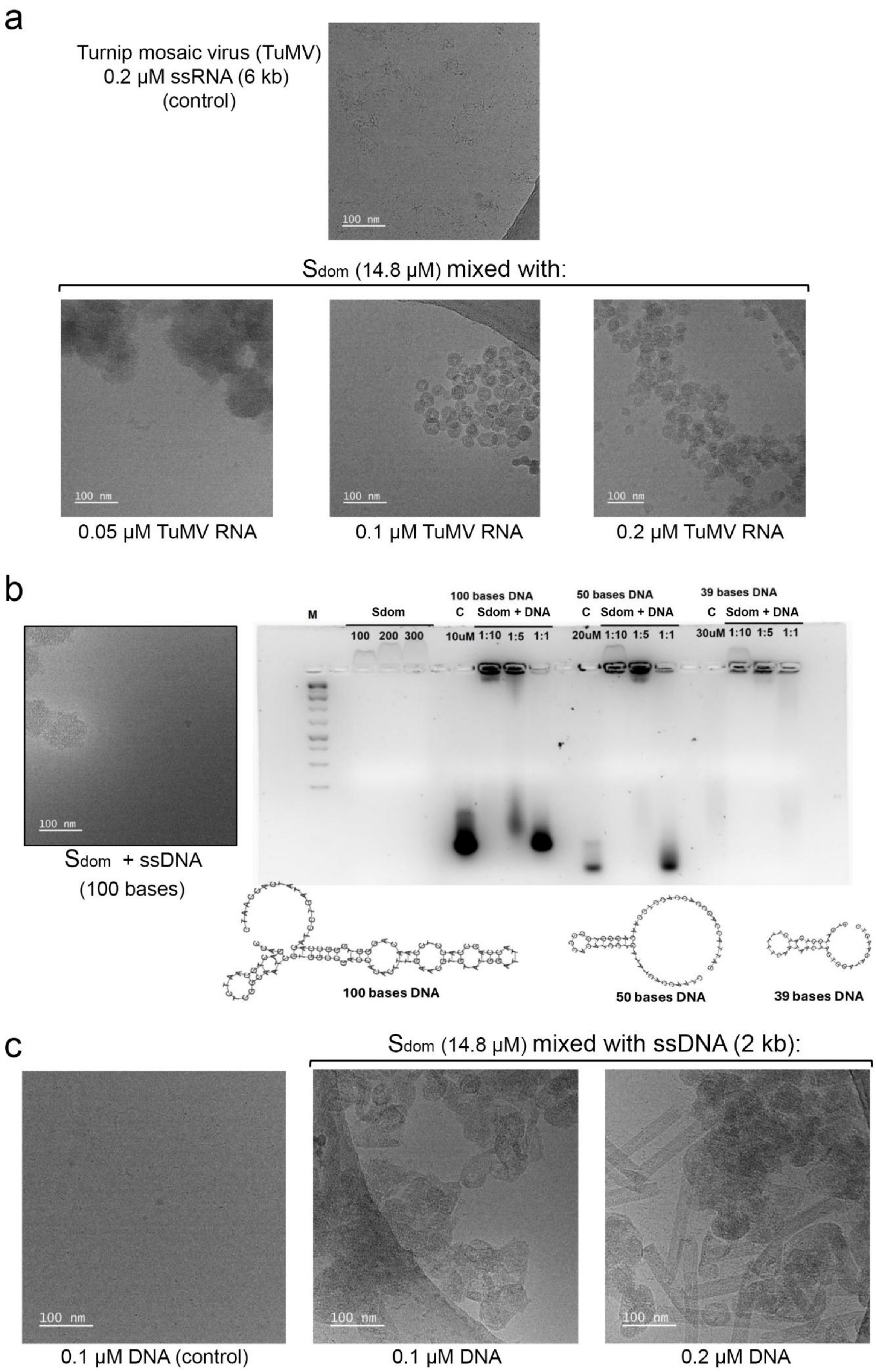
a
Turnip mosaic virus (TuMV)
0.2 µM ssRNA (6 kb)
(control)
100 nm
Sdom (14.8 µM) mixed with:
0.05 µM TuMV RNA
0.1 µM TuMV RNA
0.2 µM TuMV RNA
b
Sdom + ssDNA
(100 bases)
M
Sdom
100 200 300
100 bases DNA
C Sdom + DNA
10uM 1:10 1:5 1:1
50 bases DNA
C Sdom + DNA
20uM 1:10 1:5 1:1
39 bases DNA
C Sdom + DNA
30uM 1:10 1:5 1:1
100 bases DNA
50 bases DNA
39 bases DNA
c
Sdom (14.8 µM) mixed with ssDNA (2 kb):
0.1 µM DNA (control)
0.1 µM DNA
0.2 µM DNA

**Supplementary Figure 9 | S-domain interaction with different oligonucleotides.** (**a**) Cryo-EM micrographs showing the effect of heterologous TuMV RNA (~6 kb) on particle assembly. RNA alone (0.2 µM) does not form defined structures, whereas incubation with $S_{dom}$ protein (14.8 µM) promotes the formation of larger and more aggregated nanoparticle assemblies in an RNA concentration-dependent manner (0.05, 0.1, and 0.2 µM), indicating that assembly is largely independent of RNA sequence but influenced by RNA topology and length. (**b**) Left, representative cryo-EM micrograph of $S_{dom}$ incubated with a 100-base DNA oligonucleotide, showing poorly defined and heterogeneous assemblies. Right, EMSA analysis of $S_{dom}$ interaction with short DNA oligonucleotides of 100, 50, and 39 bases designed to adopt distinct secondary structures. Increasing oligonucleotide length and protein:nucleic acid ratio enhance retardation and high-molecular-weight complex formation, consistent with nonspecific nucleic acid binding. Schematics of predicted oligonucleotide conformations are shown below. (**c**) Cryo-EM micrographs of $S_{dom}$ incubated with longer linear DNA (~2 kb) at increasing concentrations (0.1 and 0.2 µM). DNA alone does not form defined assemblies, whereas addition of $S_{dom}$ (14.8 µM) restores higher-order organization, yielding a mixture of rounded nanoparticles and elongated tubular structures. Together, these data indicate that nucleic acid size and topology strongly modulate S-domain-driven assembly.

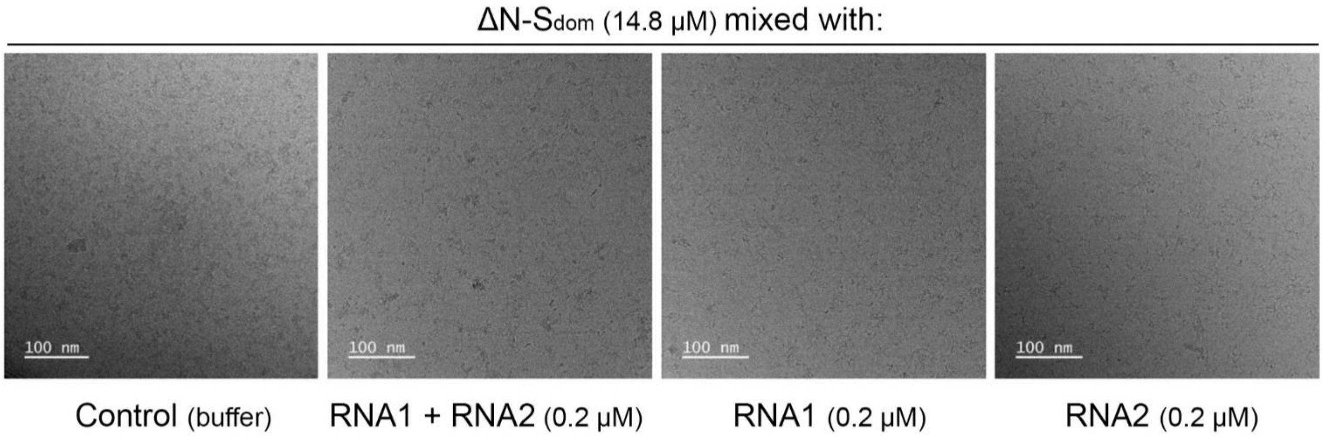


**Supplementary Figure 10 | Relevance of the N-terminal region of the S-domain in viral genome encapsidation.** Representative cryo-EM micrographs of ΔN-$S_{dom}$ (14.8 μM) incubated in buffer alone, or in the presence of RNA1+RNA2 mix (0.2 μM total), RNA1 (0.2 μM), or RNA2 (0.2 μM), showing the absence of defined nanoparticle formation under all conditions. Scale bars, 100 nm.

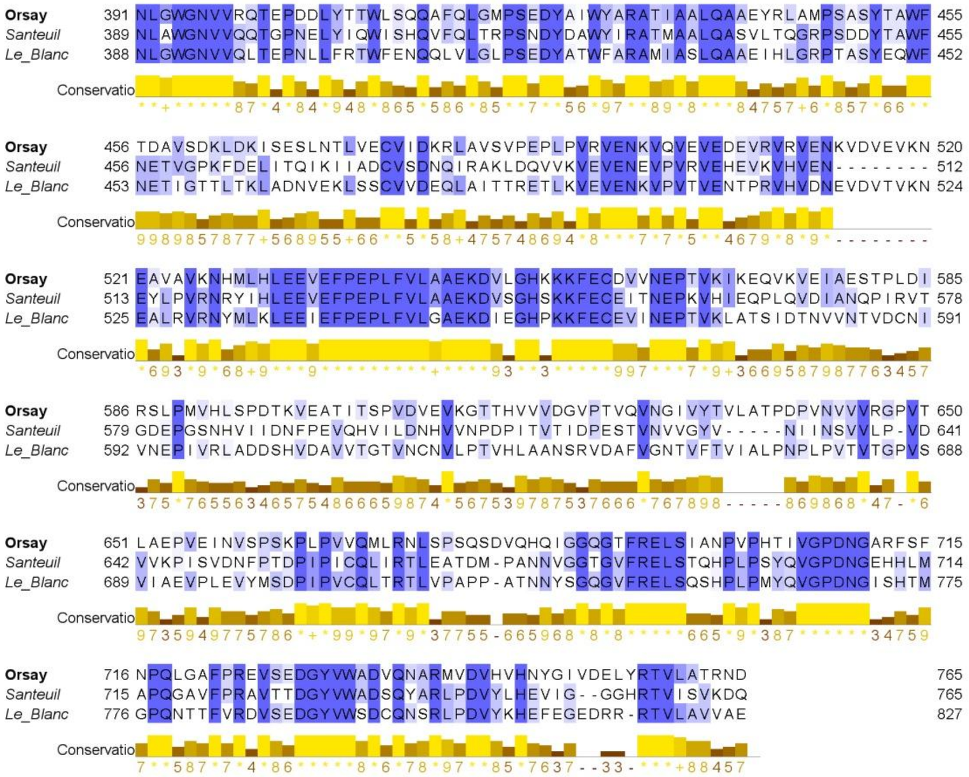

**Supplementary Figure 11 | Conservation of δ fiber anchoring motifs.** Sequence alignment of α–δ linker regions in nematode-infecting nodaviruses (OrV, Le Blanc, and Santeuil) showing conservation of residues within the linker region.

**Supplementary Table 1** | Cryo-EM data collection and refinement statistics

| **Data collection** | *Dataset A* | *Dataset A+B* |
|---|---|---|
| Microscope | Krios4 | |
| Nominal magnification | 105 kX | |
| Voltage (kV) | 300 | |
| Total dose on sample ($e^{-}$/Å$^2$) | 46.0 $e^{-}$/Å$^2$ | 50.0 $e^{-}$/Å$^2$ (*B*) |
| Frames | 50 | |
| Defocus range (µm) | -0.8 to -2.4 | |
| Sampling interval (Å/pixel) | 0.829 | |
| Nº movies | 26492 | 26492 (*A*) + 22858 (*B*) |
| **Cryo-EM processing** | *RELION - ICOS* | *cryoSPARC - ASYM* |
| Extracted particles | 60645 | 78894 |
| Contributing particles | 38101 | 77788 |
| Box Size (pixel) | 640 | |
| Pixel size (Å) | 0.829 | |
| Symmetry | I1 | C1 – local refinement (linker + δ protein) |
| Map resolution (Å)<br>FSC threshold | 2.5<br>0.143 | 2.45<br>0.143 |
| Map sharpening *B* factor (Å$^2$) | -85.37 | -94.40 |
| Deposited EMDB code | EMD-57282 | EMD-59028 |
| **Structure refinement** | *Icosahedral Asymmetric Unit* | *Peripentonal δ fragment (resid 387-452)* |
| PDB ID used as starting | 4NWV | 5JIE +AlphaFold |
| Deposited PDB ID | 29QM | |
| $CC_{mask}$ (%) | 0.86 | - |
| $CC_{vol}$ (%) | 0.75 | - |
| Model resolution (Å) | 2.5 | - |
| Model composition<br>Non-hydrogen atoms<br>Protein residues<br>Ligands ($Ca^{2+}$)<br>Water molecules | <br>16325<br>1046<br>3<br>210 | <br>2605<br>330<br>-<br>- |
| < *B* >factors (Å$^2$)<br>Protein<br>Ligands<br>Water molecules | <br>13.41<br>9.90<br>7.71 | <br>107.0<br>-<br>- |
| R.m.s. deviation<br>Bond lenghts (Å)<br>Bond angle (º) | <br>0.003<br>0.551 | <br>0.002<br>0.423 |
| Ramachandran plot<br>Favored (%)<br>Allowed (%)<br>Disallowed (%) | <br>97.79<br>2.21<br>0.0 | <br>90.62<br>9.62<br>0.0 |
| Validation Molprobity (server)<br>MolProbity score<br>All-Atoms Clashscore<br>Poor rotamers (%) | <br>0.97<br>1.99<br>0.11 | <br>1.57<br>0.99<br>1.89 |
| FSC (map, model map)<br>Threshold=0.143 (Å) | 2.3 (masked)<br>2.4 (unmasked) | -<br>- |

**Supplementary Table 2** | Primers used to generate an absolute quantification standard for RT-qPCR

| Primer name | Sequence | Application |
|---|---|---|
| oVG5_OrV_RNA1_3'_F | TAATACGACTCACTATAGGTTC CTGTCCAGGCAGTTCT | For *in vitro* transcription of OrV RNA1, and production of RNA1.3 |
| oVG6_OrV_RNA1_3'_R | GGACCTCTCCTGGGTATGTG | For *in vitro* transcription of OrV RNA1 |
| oVG7_OrV_RNA2_3'_F | TAATACGACTCACTATAGGCCT GTCAGAGTTGAGAACA | For *in vitro* transcription of OrV RNA2 |
| oVG8_OrV_RNA2_3'_R | ATAGCCGGGTATGGATAGCG | For *in vitro* transcription of OrV RNA2 |
| oVG15_RNA2_qPCR_2_F | ACGAAGCAGTAGCCGTTAAG | For RT-qPCR - at the end of OrV RNA2 |
| oVG16_RNA2_qPCR_2_R | GAGAACATCCTTCTCTGCGG | For RT-qPCR - at the end of OrV RNA2, and production of RNA2.2 |
| oVG61_OrV_RNA2_R | acggccaaagaggttatagcc | Production of RNA2 full length and RNA2.3 |
| oVG64_OrV_RNA1_R | accgtcccgtgggacctctc | Production of RNA1 full length and RNA1.3 |
| oVG226_OrV_RNA1_5'_T7_F | TAATACGACTCACTATAGGACA ATTGTCGACTTGATACCGGT | Production of RNA1 full length and RNA1.1 |
| oVG227_OrV_RNA2_5'_T7_F | TAATACGACTCACTATAGGtaaatc ggttaaccgaattacttgg | Production of RNA2 full length and RNA2.1 |
| oVG259_OrV_RNA1_R | TAGAACTGCCTGGACAGGAA | Production of RNA1.2 |
| oVG260_OrV_RNA1_T7_F | TAATACGACTCACTATAGGGGT CAAACGACCATCACACT | Production of RNA1.2 |
| oVG261_OrV_RNA1_R | AGTGTGATGGTCGTTTGACC | Production of RNA1.1 |
| oVG262_OrV_RNA2_T7_F | TAATACGACTCACTATAGGCCG CAGAGAAGGATGTTCTC | Production of RNA2.3 |
| oVG263_OrV_RNA2_T7_F | TAATACGACTCACTATAGGTTC GCTACCTTCTTCGGCTT | Production of RNA2.2 |
| oVG264_OrV_RNA2_R | AAGCCGAAGAAGGTAGCGAA | Production of RNA2.1 |
| oVGC_TuMV_T7_F | TAATACGACTCACTATAGGCTC GTTATATGGAGTCGGTTTCGG | Production of TuMV RNA |
| 25R_TuMV_R | TAACCCCTTAACGCCAAGTAAG | Production of TuMV RNA |
| RNA1 probe 1 | GCCGGGCATTTTTGGCCCTAAG TGCTTTACACTCGGACCTCGTC GACATGCATT | smFISH against RNA1 |
| RNA1 probe 2 | GGACCTGGATCGTTCTTGTTAG CCGACGTTACACTCGGACCTCG TCGACATGCATT | smFISH against RNA1 |

| | | |
|---|---|---|
| RNA1 probe 3 | CAATCTCGGGCATCTTGGTGTTAGCCAGTTACACTCGGACCTCGTCGACATGCATT | smFISH against RNA1 |
| RNA1 probe 4 | GGCGTCTGCTTTGGCCTTGCGTTCTTTTACACTCGGACCTCGTCGACATGCATT | smFISH against RNA1 |
| RNA1 probe 5 | GCTTCGATATTGAGCCTTGCGGTTTCCTTTACACTCGGACCTCGTCGACATGCATT | smFISH against RNA1 |
| RNA1 probe 6 | ACTTCGTCCATTTTGTCTTTCGTCTCGGGTTACACTCGGACCTCGTCGACATGCATT | smFISH against RNA1 |
| RNA1 probe 7 | ATGTACCACGGCATGTCGGTGCTAGTGTTACACTCGGACCTCGTCGACATGCATT | smFISH against RNA1 |
| RNA1 probe 8 | TGGTAAGCCAGTATCCGTGCGTGCTGATTACACTCGGACCTCGTCGACATGCATT | smFISH against RNA1 |
| RNA1 probe 9 | CTCGCGGATGGCATGCATCTCGTTGATTACACTCGGACCTCGTCGACATGCATT | smFISH against RNA1 |
| RNA1 probe 10 | GGCGACTGGCTCGACCTTGATCATAGTTACACTCGGACCTCGTCGACATGCATT | smFISH against RNA1 |
| RNA1 probe 11 | CGCGTTAACGGCGTTGGATCCAATAGAGCATTACACTCGGACCTCGTCGACATGCATT | smFISH against RNA1 |
| RNA1 probe 12 | CTGTCGAGCCGATAGTGAATGGCGCTCTTACACTCGGACCTCGTCGACATGCATT | smFISH against RNA1 |
| RNA1 probe 13 | CGATCGTGTATGACGTGCTGGCCTTCTTTACACTCGGACCTCGTCGACATGCATT | smFISH against RNA1 |
| RNA1 probe 14 | CCGTGGTCAAGATGATGGCGTCACCAATTTACACTCGGACCTCGTCGACATGCATT | smFISH against RNA1 |
| RNA1 probe 15 | TGCACCAACTCTTCGACGGAGTAGTGTTTCTTACACTCGGACCTCGTCGACATGCATT | smFISH against RNA1 |
| RNA1 probe 16 | GCCGGTTTTCAAGATGAACACGGGTAAGTCCTTACACTCGGACCTCGTCGACATGCATT | smFISH against RNA1 |

| | | |
|---|---|---|
| RNA1 probe 17 | AGCTACTGGAGCTGGCAGTCCGATGATTACACTCGGACCTCGTCGACATGCATT | smFISH against RNA1 |
| RNA1 probe 18 | GTATCAGAACGGAGAGCGAGTATCCCGCGTTACACTCGGACCTCGTCGACATGCATT | smFISH against RNA1 |
| RNA1 probe 19 | GCCCTAGGGGGCCCTTTGACTGTATATTACACTCGGACCTCGTCGACATGCATT | smFISH against RNA1 |
| RNA1 probe 20 | CAATTCGCTTTAGCGGTCATGCTAGCTGTTACACTCGGACCTCGTCGACATGCATT | smFISH against RNA1 |
| RNA1 probe 21 | GCAAATGTGGCTACGCTGCCGTTTACTTTACACTCGGACCTCGTCGACATGCATT | smFISH against RNA1 |
| RNA1 probe 22 | TGACGGCAGGACCTAGGTACAATTGCCTTACACTCGGACCTCGTCGACATGCATT" | smFISH against RNA1 |
| RNA2 probe 1 | "TTGGAACGCTGTCATTTTACGCCATACAGTGTTACACTCGGACCTCGTCGACATGCATT | smFISH against RNA2 |
| RNA2 probe 2 | ACAAGGCACTCCTCTGCATGTTGATGTTTACACTCGGACCTCGTCGACATGCATT | smFISH against RNA2 |
| RNA2 probe 3 | TAGACAATGCCGTTTACCTGTACCGTTGGGTTACACTCGGACCTCGTCGACATGCATT | smFISH against RNA2 |
| RNA2 probe 4 | CCTTGATCTTAACAGTTGGCTCGTTGACGACTTACACTCGGACCTCGTCGACATGCATT | smFISH against RNA2 |
| RNA2 probe 5 | ATTCGAATTTCTTCTTGTGACCGAGAACATCCTTACACTCGGACCTCGTCGACATGCATT | smFISH against RNA2 |
| RNA2 probe 6 | TGCAGCTATGGTAGCACGTGCATACCTTACACTCGGACCTCGTCGACATGCATT | smFISH against RNA2 |
| RNA2 probe 7 | GTGATATCAGGATGTGGCCCACCCTGTTACACTCGGACCTCGTCGACATGCATT | smFISH against RNA2 |
| RNA2 probe 8 | GTGTGGTCGGCCACGATTGCCGAGATTTTACACTCGGACCTCGTCGACATGCATT | smFISH against RNA2 |

| | | |
|---|---|---|
| RNA2 probe 9 | AGTCTGGTCACTTTCCTTGCGG AACGTGTTACACTCGGACCTCG TCGACATGCATT | smFISH against RNA2 |
| RNA2 probe 10 | TCAACCGTGACTGGACAGATCT GACTTGGTTACACTCGGACCTC GTCGACATGCATT | smFISH against RNA2 |
| RNA2 probe 11 | GTCGTTCGCAATGCATTGGCTT TAGCTACTTACACTCGGACCTC GTCGACATGCATT | smFISH against RNA2 |
| RNA2 probe 12 | GTTGTGTACATGGACGTCGACC ATGCGAGTTACACTCGGACCTC GTCGACATGCATT | smFISH against RNA2 |
| RNA2 probe 13 | ACTCTCGGCAATCTCGACCTTG ACTTGTTACACTCGGACCTCGT CGACATGCATT | smFISH against RNA2 |
| RNA2 probe 14 | ATCGGTTCGTCTTGGAATGAAA CCGGGACGATTACACTCGGACC TCGTCGACATGCATT | smFISH against RNA2 |
| RNA2 probe 15 | GATAGCTGAAGTGAACGGAGC CGTTACGGCTTACACTCGGACC TCGTCGACATGCATT | smFISH against RNA2 |
| RNA2 probe 16 | GGTGTCAGGTCTTTGGGAGTCT GACCGATTTACACTCGGACCTC GTCGACATGCATT | smFISH against RNA2 |
| RNA2 probe 17 | CCCAAGGCGATAGGCAATGTCT GGTGTGTTACACTCGGACCTCG TCGACATGCATT | smFISH against RNA2 |
| RNA2 probe 18 | TATGTGTGTTGTTCTTGTTTGCG TGAGGGGACTTACACTCGGACC TCGTCGACATGCATT" | smFISH against RNA2 |
